# Asymmetric Social Representations in the Prefrontal Cortex for Cooperative Behavior

**DOI:** 10.1101/2025.08.27.672249

**Authors:** Yuan Cheng, Yusi Chen, Myungji Kwak, Ross P. Kempner, Rudramani Singha, Jared Winslow, Runqi Liu, Umais Khan, Tessa Spangler, Alvi Khan, Talmo Pereira, Matthew Whiteway, Evan S. Schaffer, Nuttida Rungratsameetaweemana, Nan Yang, Herbert Zheng Wu

**Affiliations:** Department of Neuroscience and the Friedman Brain Institute, Icahn School of Medicine at Mount Sinai, New York, NY, USA; Computational Neuroscience Center, University of Washington, Seattle, WA, USA; Department of Biomedical Engineering, Columbia University, New York, NY, USA; Quantitative Sciences Unit, Department of Medicine, Stanford University, Stanford, CA, USA; Salk Institute for Biological Studies, La Jolla, CA, USA; Mortimer B. Zuckerman Mind Brain Behavior Institute, Columbia University, New York, NY, USA; Institute for Regenerative Medicine and Alper Center for Neural Development and Regeneration, Icahn School of Medicine at Mount Sinai, New York, NY, USA; Black Family Stem Cell Institute, Icahn School of Medicine at Mount Sinai, New York, NY, USA

## Abstract

Cooperation is a hallmark of social species, enabling individuals to achieve goals that are unattainable alone. Across species, cooperative behaviors are often organized by distinct social roles such as leaders and followers, yet the neural mechanisms supporting such role-based coordination remain elusive. Here we introduce a new paradigm for studying cooperation in mice, where pairs of animals engage in a joint spatial foraging task that naturally gives rise to stable leader-follower roles predictive of learning speed. Disruption of medial prefrontal cortex (mPFC) activity, particularly in followers, impairs cooperation and induces reciprocal shifts in how animals weigh self- and partner-related cues for decision-making. Calcium imaging reveals that mPFC encodes both leadership dynamics and an egocentric social value map of the partner’s position, each in an asymmetric, role-specific manner. Combining this behavior with a novel multi-agent inverse reinforcement learning framework, we identify latent value functions that guide cooperative decisions and are decodable from mPFC activity. These findings uncover fundamental neural computations that support cooperation, revealing how social roles shape decision-making in real time. Our work opens new avenues for investigating the cellular and circuit basis of social cognition and collective behavior.

## Main

Social animals learn to follow rules, adopt complementary roles, and form dynamic structures that enable effective coordination in group behavior^1,2^. A central expression of structured group behavior is cooperation, which is pervasive across species and foundational to human social life and organization^3,4^. Among the most common strategies for organizing cooperative behavior are leadership and followership^5^, roles that allow individuals to influence and respond to one another in pursuit of common goals. The leader-follower dynamics are ubiquitous across taxa, from the flight of birds^6^ to human institutions^7,8^. Despite their importance, the cellular and circuit-level mechanisms of how social roles support cooperation remain largely unknown. In particular, a robust and ethologically grounded paradigm for studying cooperation is still lacking in mice, posing a critical barrier to harnessing the modern neuroscience tools in this model organism^9,10^. Consequently, a comprehensive mechanistic understanding of cooperative behavior remains elusive.

A central question in the study of cooperation is how individuals make social decisions^11^. In natural settings, social cues are dynamic and actions unfold continuously rather than in discrete, turn-based games^12–14^, requiring moment-to-moment coordination^15^. Further, social decisions are likely shaped not only by immediate sensory^16–20^ or value-based processes^21–25^, but also by experience, identity, and role^26–30^. Another key challenge in studying complex social interactions is that an agent’s underlying goals and strategies are often unobservable, especially when multiple agents interact reciprocally. Although inverse reinforcement learning (IRL), a form of learning from observation, offers a principled way to infer an agent’s internal model of the world from observed behavior^31,32^, extending IRL to multi-agent biological systems introduces significant complexity. We still lack a mechanistic understanding of how individual goals contribute to joint decision-making and how this process is shaped by social roles like leader and follower.

To address these questions, we developed a novel cooperative foraging paradigm in mice and a multi-agent inverse reinforcement learning (MAIRL) framework. We found that the medial prefrontal cortex encodes leadership dynamics and an egocentric “social value map” of partner’s position in a role-specific and task-dependent manner, mirroring the latent goals of the animals uncovered by MAIRL. These findings uncover the neural computations underlying role-based social decision-making in cooperation, offering cellular-resolution insights into the neurobiological basis of cooperative behavior.

### Robust leader and follower roles emerge in cooperative foraging

In the cooperative foraging paradigm, pairs of water-restricted, same-sex adult mice freely interact in a square open arena to obtain water rewards (Fig. 1a). Each arena wall features a reward zone with two ports, enabling simultaneous access. A trial begins when either mouse touches a central initiation spot, triggering the random activation of two of the four reward zones, indicated by LED lights. To earn a reward, both mice must navigate to the same active zone and make concurrent nose pokes into the two ports. We define “cooperation” here as joint action of two agents toward a shared goal, namely, obtaining water through spatiotemporal coordination. Trials end in error if mice poke concurrently in different active zones (“mismatch error”) or either mouse pokes in an inactive zone (“unrewarded error”), resulting in no reward and a brief timeout (Methods). The vast majority of the pairs reached the training criterion—80% correct trials across three consecutive sessions—with females learning faster and performing better at criterion than males (median: 11 vs. 19 days; interquartile range: 9–19 vs. 12–24; Fig. 1b, Supplementary Fig. 1a-f, Supplementary Video 1). Interestingly, with training, mouse pairs significantly improved the synchronization of their nose pokes at the reward ports, even though precise timing was not required, as one mouse could wait for the other by holding its nose poke (Supplementary Fig. 1g). We used machine learning tools, SLEAP^33^ and EKS^34^, to annotate the nose, neck, and torso positions of both mice in each video frame, enabling precise tracking of their movement and posture.

**Figure 1.**
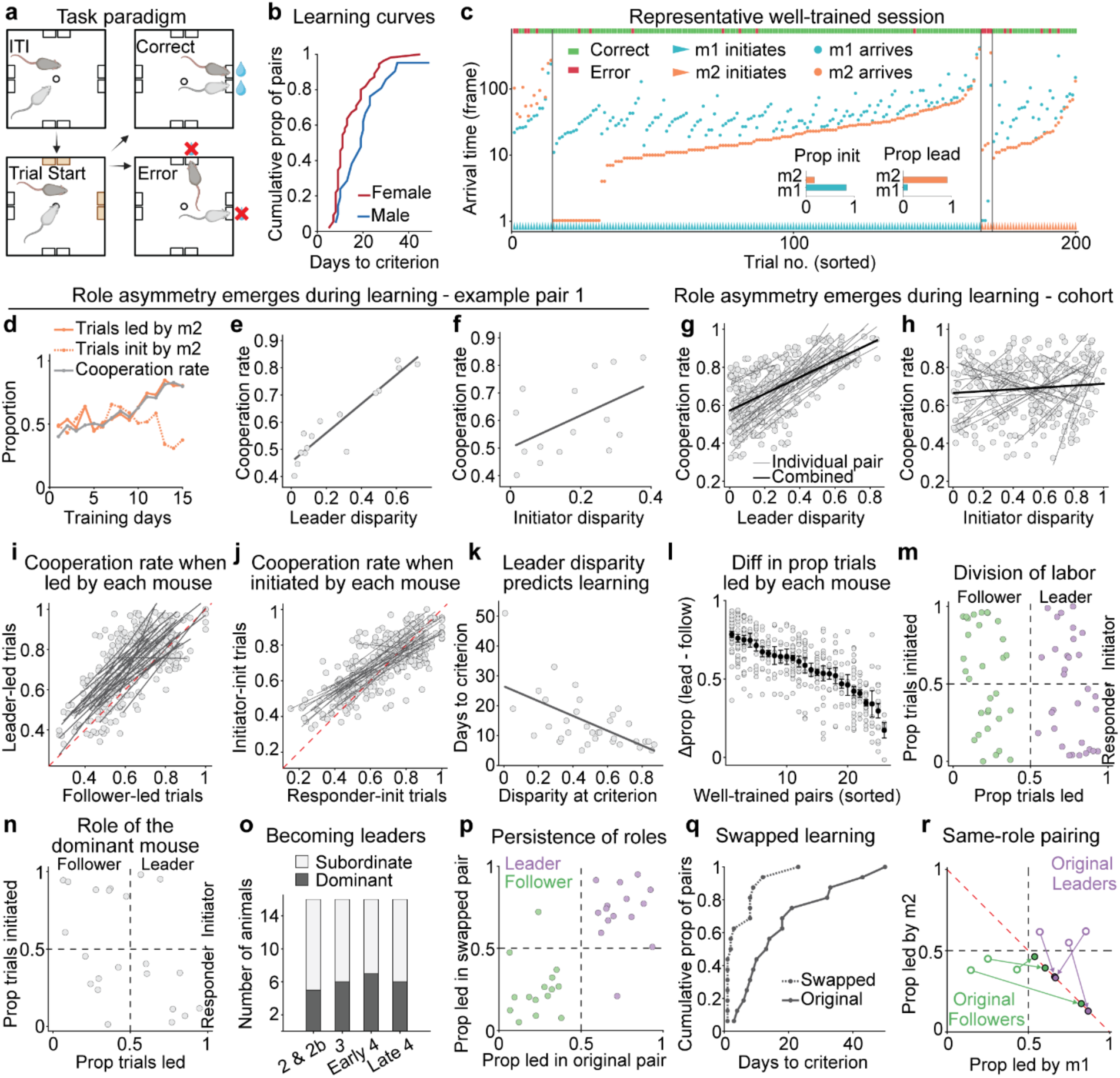
Cooperative foraging gives rise to stable social roles. **a**, Schematic of the cooperative foraging paradigm. Created in BioRender. Wu, H. (2025) https://BioRender.com/8sa1c1v. Same below. **b**, Cumulative proportion of mouse pairs reaching the training criterion over days in the final training stage. Female pairs outperform males (two-tailed Kolmogorov-Smirnov test, *P* = 0.036; *n* = 55 female, 21 male pairs). **c**, Representative well-trained session showing trial-by-trial performance. Each column represents a single trial, sorted by arrival time, and grouped by initiator and leader identity. Trial outcomes are indicated by colored bars at the top (green: correct; red: error). Triangles at the bottom denote the initiating mouse (mouse #1: cyan; mouse #2: orange), and circles indicate arrival times. Video recorded at 30 frames per second. Insets show the proportion of trials initiated and led by each mouse: m1 initiated 83% of the trials, while m2 led 91%. **d,** Cooperation rate alongside the proportion of trials led (solid curve) or initiated (dashed curve) by one mouse in an example pair across days. Data for the partner mouse are not shown, as these values are complementary. **e,** Correlation between cooperation rate and leader disparity (absolute difference in the proportion of trials led by each mouse in a pair). Pearson’s correlation: *R* = 0.93, *P* = 3.5×10^-7^, *n* = 15 sessions. **f,** Correlation between cooperation rate and initiator disparity (absolute difference in the proportion of trials initiated by each mouse in a pair). Pearson’s correlation: *R* = 0.50, *P* = 0.056. **g,h,** Correlation between role asymmetry and cooperation rate during learning across pairs. Each point represents one session from a mouse pair. Thin lines show linear fits for individual pairs; thick line for all sessions combined. Leader disparity is strongly correlated with cooperation rate (**g**, linear mixed-effects model, *P* = 2.5×10^-54^; *n* = 24 pairs, 306 sessions), while initiator disparity shows a weak correlation with greater variability across pairs (**h**, *P* = 0.038). **i,j,** Cooperation rate in trials led (**i**) or initiated (**j**) by each mouse. Each point represents one session from a mouse pair. Gray lines show linear fits for individual pairs; red dashed line indicates unity. Performance is significantly higher in trials led by leaders (**i**; linear mixed-effects model, *P* = 4.4 × 10^-10^; *n* = 24 pairs, 306 sessions) and initiated by initiators (**j**; *P* = 3.4 × 10^-5^). **k**, Leader disparity at criterion significantly correlates with training duration. Each point represents one mouse pair. Pearson’s correlation, *R* = -0.60, *P* = 1.5x10^-4^; *n* = 34 pairs. **l**, Difference in proportions of trials led by leader and follower across sessions from well-trained pairs. Open circles represent individual sessions; filled circles and error bars indicate the mean ± SEM for each pair. The difference is significantly different from chance for every pair. Binomial test against 0.5, *P* < 4.3x10^-9^ for all 26 pairs; *n* = 5-32 sessions and 696-4398 trials per pair. **m**, Leaders and followers can adopt either the initiator or responder role across pairs. The proportion of trials initiated by the leader does not differ significantly from chance (two-tailed Wilcoxon signed-rank test against 0.5, *P* = 0.56; *n* = 31 pairs). Only pairs with a statistically significant leader at the training criterion (binomial test against 0.5, *P* < 0.01) are included. **n**, The proportion of trials initiated or led by the dominant animal of a pair does not differ from chance (two-tailed Wilcoxon signed-rank test against 0.5, *P* = 0.98 for initiation, *P* = 0.51 for leadership; *n* = 23 pairs). All initiation/leadership combinations also do not differ from chance (binomial test against 0.25; all *P* > 0.098). **o**, Number of dominant and subordinate mice identified as leaders in the same group of animals across different training stages. Proportions are not significantly different from chance (two-tailed binomial test against 0.5, *P* > 0.21 for all four stages). No significant differences in proportions across stages (χ² (3, *n* = 64) = 0.52, *P* = 0.91). **p**, Most leaders and followers (13 out of 15) maintain their roles in the swapped pairs (one-tailed binomial test against 0.5, *P* = 0.0037). **q**, Cumulative proportion of mouse pairs reaching training criterion over days, comparing learning in the original vs. swapped pairings (two-tailed Kolmogorov-Smirnov test, *P* = 0.0071; *n* = 10 female pairs and 6 male pairs). **r**, Change in leader-follower dynamics when pairing two original leaders or two original followers. Each arrow represents a swapped pair. Open circle indicates the proportions led by each mouse in their respective original pairs; closed circle, the newly formed pair. Convergence on the diagonal indicates role adjustment to re-establish new leaders and followers (binomial test against 0.5, *P* < 0.019 for all 6 new pairs; 304-1,049 trials per pair).

Consistent behavioral roles emerged as mice learn to cooperate (Fig. 1c). Once well-trained, one mouse typically initiates more trials and is termed the initiator, while the other becomes the responder. Separately, one animal consistently arrives first at the reward zone and is defined as the leader, with the other as the follower (Methods). We use “leader” here in its literal sense: one who guides by going in advance. Strikingly, role differentiation appeared to facilitate learning: the earlier these roles emerge, the faster the pair learns to cooperate (Fig. 1d-h, Supplementary Fig. 1h-j). Cooperation rate, measured as the proportion of match trials over total match and mismatch trials (Methods), was higher in trials led by the leader (Fig. 1i) and initiated by the initiator (Fig. 1j), indicating a behavioral advantage from role adherence. Mice with weaker leader-follower disparity still learned the task but required more time (Fig. 1k). Once established, these roles remained stable across sessions (Fig. 1l). Although initiators appeared less likely to lead in some pairs (Fig. 1c,d, Supplementary Fig. 1h), this division of labor was not significant overall, and individual mice can assume both initiator and leader roles (Fig. 1m). Given the role of dominance hierarchy in many forms of mouse social behavior, particularly competitive interactions^35–37^, we tested whether social roles in the cooperative task might also reflect dominance status, determined by the tube test^38^. Surprisingly, neither role correlates with dominance at well-trained or earlier stages (Fig. 1n,o, Supplementary Fig. 1k), indicating that these roles constitute distinct social structures independent of hierarchy.

Because social roles can persist across contexts and partners, we next asked whether mice maintain their established roles when paired with new individuals. We swapped two well-trained pairs such that the leader of one pair was teamed with the follower of another. Most animals maintained their original roles in the new pairings, demonstrating that leader-follower identity is a stable construct carried across contexts (Fig. 1p). Interestingly, while some new pairs reached the training criterion immediately after swapping, others required time to adapt—though still less than during initial learning (Fig. 1q). When two leaders or two followers were paired, one mouse adopted the complementary role to enable cooperation (Fig. 1r). These results underscore the inherently social nature of the task: mice preserve established roles even with unfamiliar partners, yet can flexibly adopt new ones when needed, with variable adaptation times reflecting the need to calibrate with different individuals.

Well-trained mice adopted a set of stereotyped movement patterns, or “behavioral motifs”, to solve the task (Fig. 2a, Supplementary Fig. 2a, Supplementary Videos 2-5). We defined four motifs based on empirical observation and manually annotated them (Methods). All the motifs involve pair-level interaction, but “track”, “sharp turn”, and “join” were annotated with respect to the actor mouse, while “synchronized travel (sync)” was not assigned to a specific individual, as it reflects a joint behavior. In the “track” motif, for example, one mouse slows down near the reward zone, turns to monitor its partner’s movement, and allows the partner to catch up to align the arrival times. In “sharp turn”, one mouse abruptly redirects its path to follow its partner. These motifs may reflect varying degrees of coordination between the animals. Moreover, animal pairs exhibited systematic differences in motif profiles, indicating distinct individual strategies (Fig. 2b-d). Leaders primarily use the “join” motif, especially when they follow (Fig. 2e), while followers show more varied motifs and increase “sharp turns” when required to follow more often (Fig. 2f). These findings suggest that mouse pairs develop individual, role-specific strategies and flexibly adapt to changing demands for cooperation.

**Figure 2.**
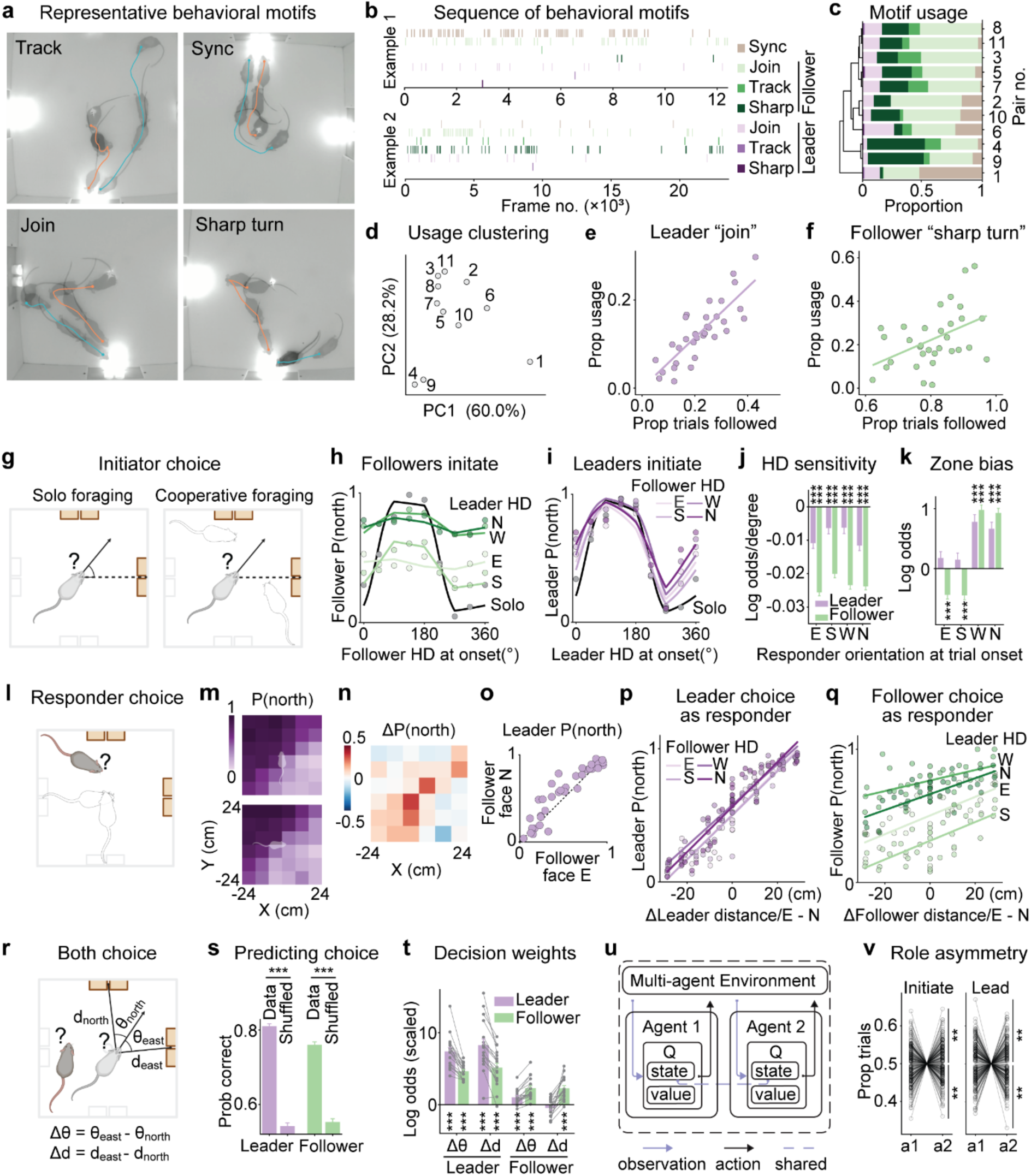
Social roles shape decision-making through asymmetric yet reciprocal interactions. **a**, Representative behavioral motifs. Cyan and orange curves indicate mouse trajectories. Each motif is illustrated by overlaying 3-4 video frames. **b**, Gantt plots of motif sequences from two example sessions, showing only concatenated within-trial frames. **c**, Motif usage across 11 pairs, averaged over 2-4 sessions per pair, with hierarchical clustering based on similarity in motif composition. Color scheme as in **b**. **d**, Principal component analysis (PCA) of motif usage across pairs. The first two PCs account for 60.0% and 28.2% of the total variance, respectively. **e**, Usage of “join” in leaders correlates with the proportion of trials they follow. Each point represents one session. Pearson’s correlation: *R* = 0.83, *P* = 6.1 × 10^-9^; *n* = 32 sessions, 12 pairs. **f**, Usage of “sharp turn” in followers correlates with the proportion of trials they follow. Pearson’s correlation: *R* = 0.45, *P* = 0.0093; *n* = 32 sessions, 12 pairs. **g**, Schematic for comparing initiator decisions during solo versus cooperative foraging. Trials with adjacent active zones are rotated to the north-east orientation. Heading is defined as the angle between the neck-nose vector and the x-axis. **h**, Follower choice for the north zone, P(north), plotted against its heading at trial onset. Each point indicates the proportion of trials in which the follower selected north, binned in 45° intervals centered on the x-axis values. Curves show logistic fits of P(north) as a function of follower heading, conditioned on leader’s heading at trial onset (facing east, south, west, or north), or during solo foraging sessions. **i**, Leader choice plotted against heading. Format as in **h**. **j,k**, Beta estimates from logistic regression showing how responder orientation affects initiator’s sensitivity to its own heading and initiator’s bias in choice (positive values indicate a bias toward the north; negative, east). **j**, When leader is the responder, its presence significantly reduces follower’s sensitivity (logistic mixed-effects regression, all \*\*\**P* < 7.8 ×10^-72^; *n* = 8 pairs and 4,991-5,129 trials per condition) and vice versa (all \*\*\**P* < 4.5 × 10^-4^; *n* = 8 pairs, 3,881-3,963 trials per condition). **k**, Leader facing east and south decreases follower’s choice bias, while west and north increases it (all \*\*\**P* < 1.2 ×10^-9^). Follower facing west and north significantly increases the leader bias (both \*\*\**P* < 1.5 × 10^-9^). **l**, Schematic for comparing responder choices across initial position, conditioned on initiator orientation at trial onset. **m**, Leader choice P(north) across initial positions at trial onset as a responder, when the follower is facing north (top) or east (bottom). The cartoon mouse indicates the orientation of the follower as the initiator. **n**, Leader ΔP(north), calculated as P(north | follower facing north) - P(north | follower facing east). **o**, Leaders are more likely to choose north when the followers face north. Each point denotes a spatial bin from the left plots. Paired *t*-test, *P* = 0.0049; *n* = 36 bins, 4,433 trials (follower facing east) and 4,464 trials (follower facing north) pooled across 16 pairs. **p**, Leader choice plotted against the difference in its distance to the east and north zones, conditioned on follower orientation at trial onset. Each point is the P(north) within a spatial bin; curves show linear fits. P(north) is significantly higher when the follower faces west or north than east or south (repeated measures ANOVA across four conditions, followed by Tukey-Kramer post hoc test, all *P* < 0.025 for east/south vs. west/north, *P* > 0.88 for both east vs. south and west vs. north). **q**, Follower choice plotted against difference in distance to the east and north zones, conditioned on leader orientation at trial onset. P(north) ranked highest to lowest when the leader faced west, north, east, and south (all pairwise comparisons *P* < 1.1 ×10^-5^). **r**, Schematic for examining both initiator and responder choices across initial conditions. For each animal, we quantified the signed difference in head angle (Δθ) relative to the east and north zones (θ_east_ and θ_north_), and the signed difference in distance (Δd) to these zones (d_east_ and d_north_). **s**, Logistic regression using Δθ and Δd from both animals (scaled to [0, 1] range) to predict the leader or follower choice. Cross-validated predictions significantly outperform shuffled controls (bootstrap test, *P* < 1.0 x 10^-4^, 10,000 iterations). **t**, Beta coefficients for predicting leader or follower choice. Each point represents an individual animal, and lines connect animals from the same pair. Bars and error bars indicate the mean ± SEM. All groups are significantly different from zero except follower’s distance weight (Δd) for leader choice (one-sample *t*-test, \*\*\**P* < 9.0 x 10^-4^; *n* = 17 pairs, 668-3,235 trials per pair). Coefficients also differ significantly between leaders and followers (paired *t*-test; all *P* < 3.3 x 10^-5^). **u**, Schematic of the two-agent decision-making simulation using MARL. Each agent observes its own state, the partner’s state, and active reward zone locations, and selects actions through a separate Q module mapping value to state. Agents share a common state representation, denoted by the dashed line. **v**, Asymmetry in adopting the initiator (left) and leader (right) roles. Circles with connecting lines show proportions of trials initiated or led by agent 1 versus agent 2 for pairs with significant bias (binomial test, *P* < 0.01); grey lines without circles indicate pairs without significant bias.

### Social roles shape decision-making through asymmetric yet reciprocal interactions

We next examined how mice make foraging decisions based on their leader or follower roles. We focused first on trials when they are the initiators, comparing their reward zone choices in cooperation to a solo foraging control (Fig. 2g). In this control, mice foraged alone in the absence of partner influence and were rewarded at either of the two active zones. For analysis, we rotated trial types with adjacent active zones to the north and east (Methods). We then asked how an initiator’s trial-by-trial choice (north vs. east) depends on their heading, conditioned on the responder’s heading or position at trial onset.

In solo foraging, both leaders and followers showed clear heading-dependent choices, e.g., animals facing north at trial onset were more likely to choose the north zone (Fig. 2h,i, black curves). In cooperative foraging, this heading dependence was preserved but was strongly modulated by the responder’s heading and position. Followers, in particular, showed reduced sensitivity to their own heading and increased bias toward the leader’s heading (Fig. 2h,j,k). This influence was reciprocal: leaders were also influenced by follower’s heading, showing modest but still significant reductions in sensitivity and corresponding behavioral biases (Fig. 2i-k). Similar effects were observed when considering the partner’s position (Supplementary Fig. 2b-e). These results suggest that both roles flexibly integrate partner’s positional information into decision-making, with followers showing greater sensitivity to leader’s state.

We then analyzed trials in which leaders or followers were the responders, who could be at any location in the arena at trial onset (Fig. 2l). After rotating the active zones to the east and north, leaders exhibited a clear spatial gradient in choice preference, favoring the closer reward zone (Fig. 2m). Notably, this preference was strongly influenced by follower’s heading: when followers faced north at trial onset, leaders were more likely to choose north, particularly at equal distance to east and north zones (Fig. 2n-o). By analyzing leader choices as a function of their relative distance to the two active zones, we found that follower heading consistently modulated leader decisions across all directions (Fig. 2p). Similarly, leaders influenced follower choices (Fig. 2q). These results highlight the reciprocal influence in decision-making by each animal’s head orientation.

Finally, we used logistic regression to fit the decisions of both leaders and followers across all trials using four predictors: each animal’s position and heading difference relative to the two active zones at trial onset (Fig. 2r). This simple regression reliably predicted zone choice (north vs. east) for both animals, validating its effectiveness (Fig. 2s). Notably, leader’s heading and position exerted a stronger influence on follower’s choice than the follower’s own spatial variables (Fig. 2t). Conversely, the follower’s heading, but not position, significantly impacted the leader’s decision. These results corroborate the asymmetric yet reciprocal influence between the two roles during cooperative foraging.

### Spontaneous role differentiation in simulated cooperative behavior

To examine how coordinated strategies and asymmetrical roles might arise from shared incentives alone, we first modeled cooperative behavior using forward multi-agent reinforcement learning (a data-driven inverse modeling follows later in the manuscript). In this framework, agents learn through experience to choose actions that maximize rewards over time by updating a value function, which estimates how rewarding each action is. This approach allows us to test whether role differentiation can emerge without prior biases or explicit role assignments. Our simulation mirrors the mouse task design, and the training used a two-stage curriculum mirroring mouse training (Methods). Each agent independently learned a value function based on the joint spatial state using one-step temporal difference learning (Fig. 2u).

All simulated agent pairs learned the task, achieving accuracy comparable to or exceeding that of mice (Supplementary Fig. 2f-h). Despite identical initializations, behavioral asymmetries reliably emerged across pairs (Supplementary Fig. 2i). During testing, 183 pairs established statistically significant initiators and 183 consistent leaders (Fig. 2v). As in mice, fewer initiators were also leaders, but this difference was not significant, suggesting that initiation and leadership are dissociable roles (Supplementary Fig. 2j). These results suggest that asymmetric social roles can emerge spontaneously, driven by small biases in agents’ learned value maps that favor certain spatial configurations of the pair over their mirrored counterparts. These biases likely originate from stochastic fluctuations during learning and are amplified and stabilized by coordination demands. This echoes mouse behavior, where stable social roles emerge among genetically identical animals raised in the same environment. The simulation thus offers a computational account of how social roles can self-organize through interaction alone, without preexisting individual differences.

### Role-specific dependence of cooperative behavior on mPFC

We next sought to identify the brain areas necessary for cooperative behavior in mice. The prefrontal network plays an important role in social cognitive processing in humans and primates^39,40^. In rodents, the medial prefrontal cortex (mPFC) has been implicated in sociality^41,42^, dominance^38^, interbrain synchrony^43^, and competitive behavior^36,37^, and is critical for transforming accumulated evidence into discrete choices during goal-directed solo tasks^17^. The orbitofrontal cortex (OFC) plays key roles in representing value^44^ and task space^45^. While both mPFC and OFC have been shown to encode cooperation-related variables in rodents^46^, their causal role in cooperative behavior remains unknown.

We examined the causal roles of mPFC and OFC in cooperative behavior using the inhibitory DREADD receptor hM4D(Gi) and validated its efficacy in awake animals via *in vivo* extracellular recordings (Supplementary Fig. 3a,b). Bilateral inactivation of mPFC, but not OFC, significantly reduced cooperation rates (Fig. 3a,b; Supplementary Fig. 3c,d). By contrast, performance remained intact in mCherry-expressing controls (Supplementary Fig. 3e,f). The performance decline was driven by a significant increase in mismatch errors, rather than unrewarded errors, indicating a specific impairment in coordination rather than general deficits in reward-seeking or navigation (Supplementary Fig. 4a,b). mPFC inactivation also diminished leader disparity (Fig. 3c) without affecting initiator disparity (Fig. 3d), indicating a disruption of leader-follower dynamics. Cooperation declined on both leader-and follower-led trials (Fig. 3e,f), suggesting a general impairment in coordination. Notably, only followers showed a significant reduction in reaction time (Fig. 3g,h), possibly reflecting less deliberation. These effects cannot be attributed to an impairment in locomotion (Supplementary Fig. 4b,c). Logistic regression revealed that followers placed greater decision weights on their own heading and less on leader position, whereas leader strategies remained unchanged (Fig. 3i).

**Figure 3.**
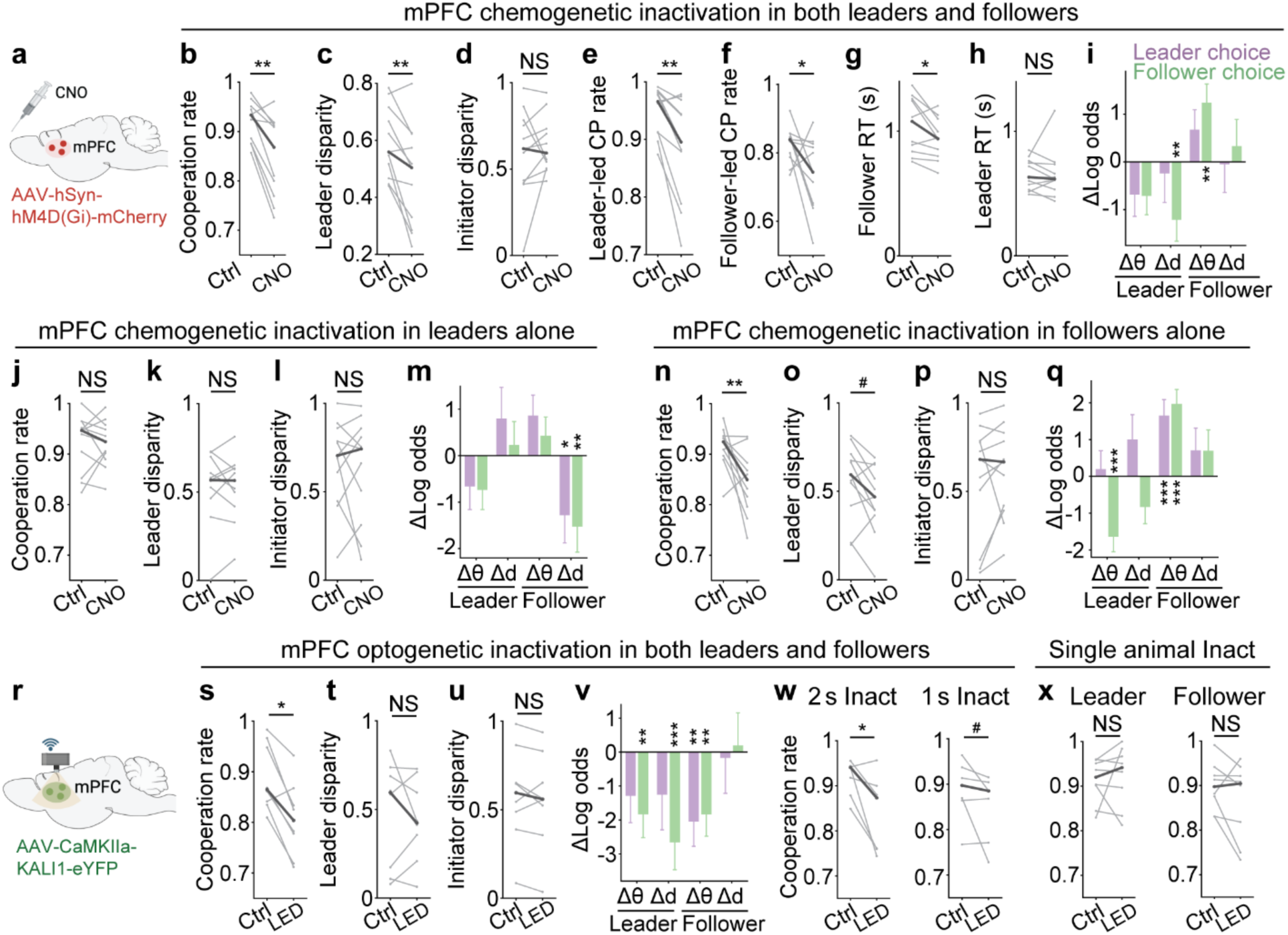
Role-specific dependence of cooperative behavior on mPFC. **a**, Schematic of chemogenetic inactivation in the mPFC. **b-h**, mPFC inactivation in both leaders and followers significantly reduces the cooperation rate (two-tailed Wilcoxon signed-rank test, *\**\**P* = 0.0029; *n* = 11 pairs), leader disparity (*\**\**P* = 0.0068), and cooperation rate (cp rate) in trials led by the leader (*\**\**P* = 0.0049) or by the follower (*\*P* = 0.042). It does not affect initiator disparity (*P* = 0.32; NS, not significant). Mean reaction time (in sec) is significantly reduced in followers (\**P* = 0.024), but not in leaders (*P* = 0.41). **i**, Changes in beta coefficients from logistic regression for predicting leader or follower choice. Bars show estimate ± standard error. mPFC inactivation significantly reduced weighting for leader’s distance (Δd) (\*\**P* = 0.0076; *n* = 11 pairs, 6,798 trials) and increased weighting for follower’s heading (Δθ) (\*\**P* = 0.0014) in follower decisions. **j-l**, mPFC inactivation in leaders alone did not affect cooperation rate, leader disparity, or initiator disparity (two-tailed Wilcoxon signed-rank test, all *P* > 0.76, *n* = 11 animals). **m**, Leader inactivation alone significantly reduced the weighting for follower’s distance in leader and (logistic regression, \**P* = 0.031; *n* = 6,691 trials) follower decisions (\*\**P* = 0.0053). **n-p**, mPFC inactivation in followers alone reduced the cooperation rate (two-tailed Wilcoxon signed-rank test, *\**\**P* = 0.0098; *n* = 11 animals) and showed a trend toward decreased leader disparity (*^#^P* = 0.054), with no significant effect on initiator disparity (*P* = 0.17). **q**, Follower inactivation alone significantly reduced the weighting for leader’s heading (logistic regression, *\**\*\**P* = 4.5 x 10^-5^, *n* = 6,805 trials) and increased the weighting for follower’s own heading in follower decision (*\**\*\**P* = 5.2 x 10^-7^). The weighting for follower’s heading shows a significant increase in leader decision (*\**\*\**P* = 1.4 x 10^-4^). **r**, Schematic of wireless mPFC optogenetic inactivation. **s-u**, Optogenetic inactivation in both leaders and followers for the entire trial duration impaired the cooperation rate (two-tailed Wilcoxon signed-rank test, \**P* = 0.016; *n* = 8 pairs) but not leader disparity (*P* = 0.74) or initiator disparity (*P* = 0.46). **v**, Inactivation significantly reduced the weighting for follower’s heading in leader decision (logistic regression, \*\**P* = 0.0050; *n* = 1,188 trials) and reduced the weightings for leader’s heading (\*\**P* = 0.0074), leader’s distance (*P* = 9.3 x 10^-4^), and follower’s heading (\*\**P* = 0.0041) in follower decision. **w,** Inactivation for 2 s (left) from trial onset significantly impaired cooperation rate (two-tailed Wilcoxon signed-rank test, \**P* = 0.031, *n* = 7 pairs), while 1 s inactivation (right) showed a trend toward impairment (^#^*P* = 0.078). **x**, Inactivation in either leaders (left) or followers (right) alone for the trial duration did not significantly affect cooperation rate (two-tailed Wilcoxon signed-rank test, *P* = 0.95 and *P* = 0.11 respectively; *n* = 8 pairs).

We next asked how each social role could contribute to the joint decision-making by selectively inactivating mPFC in either the leader or the follower. Inactivating the mPFC in leaders did not significantly impair task performance or disrupt the role dynamics (Fig. 3j-l). Leaders showed reduced weighting of follower distance in decision-making, which was paralleled by a similar reduction in how non-inactivated followers weighted their own distance (Fig. 3m), possibly indicating a compensatory adjustment. In contrast, inactivating follower mPFC reduced cooperation rates without significantly altering the role dynamics (Fig. 3n-p). Logistic regression showed that inactivated followers relied more on their own heading and less on partner cues (Fig. 3q), echoing patterns observed during the inactivation of both roles. Interestingly, the non-inactivated leaders increased sensitivity to the follower’s heading, suggesting they may also compensate for impaired follower performance to maintain coordination.

Building on our chemogenetic findings, we next applied wireless optogenetics to achieve greater temporal precision in mPFC silencing during free interactions (Fig. 3r), validated via *in vivo* recordings (Supplementary Fig. 4d-g). Trials were randomly assigned to receive inactivation from onset to offset, with the remainder serving as within-session controls (Methods). Optogenetic inactivation impaired task performance (Fig. 3s), recapitulating the main effect seen with chemogenetic silencing, but did not disrupt leadership or initiatorship (Fig. 3t,u). Both roles showed reduced weighting of position and heading cues during decision-making (Fig. 3v). Notably, brief inactivation for just 1-2 s from trial onset was sufficient to impair performance, highlighting a critical early decision period (Fig. 3w). However, silencing either the leader or follower alone did not impair behavior (Fig. 3x), suggesting that intact mPFC activity in at least one animal can support coordination. These results confirm that mPFC is essential for cooperation and highlight the reciprocal nature of this interaction, which provides resilience against transient disruption in either partner.

### Asymmetric representations of leadership dynamics in mPFC

Having established a causal role for mPFC in cooperative behavior, we next asked how social roles and decisions are represented at the neural population level. We recorded mPFC activity using one-photon calcium imaging in freely interacting mouse pairs performing the foraging task. As expected, many mPFC neurons encoded task-relevant signals such as choice (Fig. 4a). Interestingly, a greater proportion of neurons were selective for choice in leaders compared to followers (Fig. 4b). To evaluate the population-level representation, we trained decoders to predict the reward zone choices from mPFC activity. Decoding accuracy was consistently higher in leaders, suggesting that mPFC representations of behavioral choices are more robust in leadership roles (Fig. 4c).

**Figure 4.**
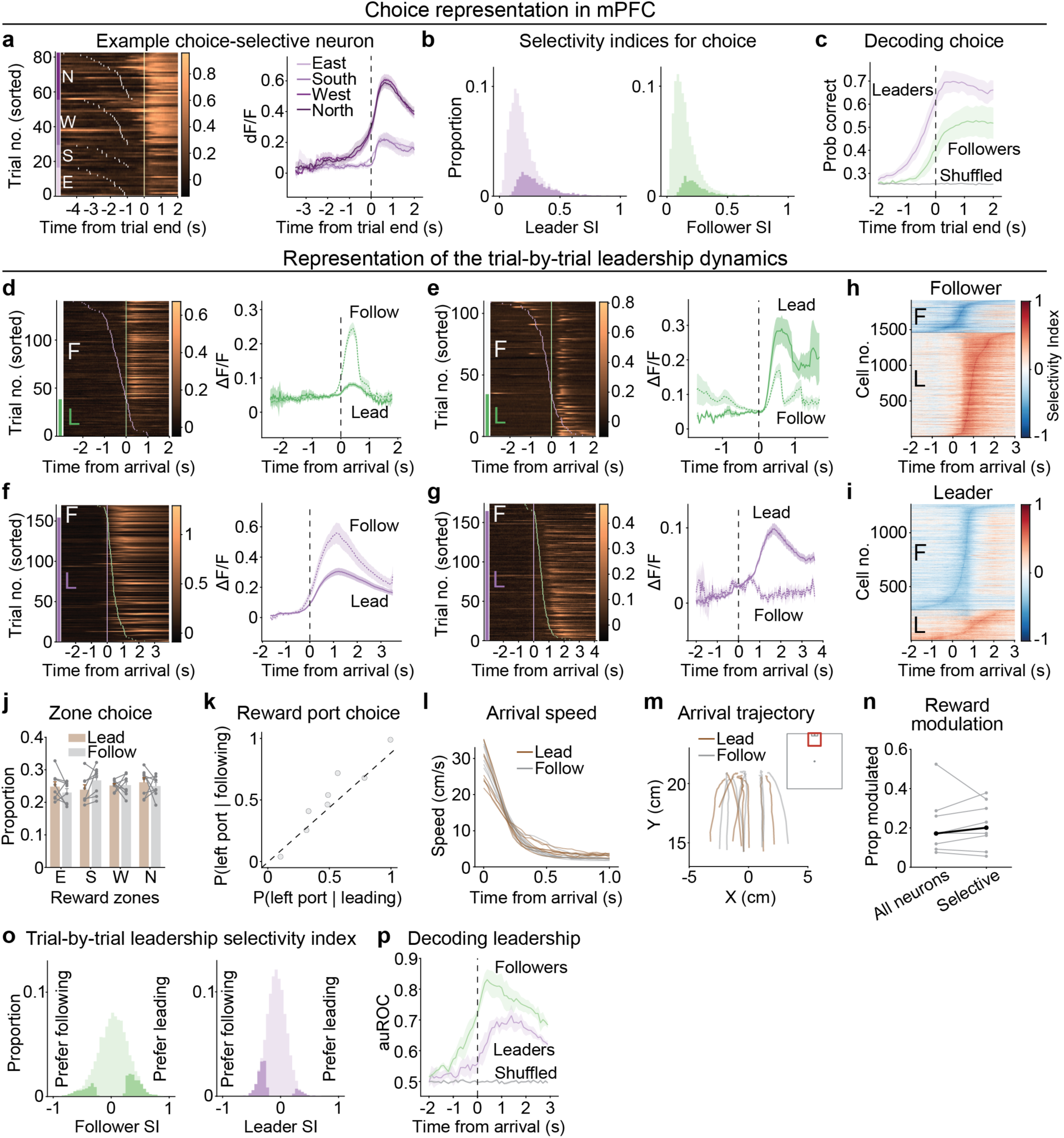
Asymmetric representations of leadership dynamics in mPFC. **a**, Example choice-selective neuron recorded from a leader mouse. Left: trial-by-trial calcium activity (ΔF/F). Each row is a trial, grouped by the reward zone chosen by the mouse and sorted by trial duration. The white and yellow ticks indicate the onset and offset of the trials, respectively. Right: Curves and shading show mean ΔF/F ± SEM across trials. **b**, Selectivity indices (SIs) for choice during the 1-s time window before trial end. Left: leaders (*n* = 8,395 neurons, 4 animals); right: followers (*n* = 6,995 neurons, 4 animals). Shaded areas indicate neurons with significant selectivity (one-way ANOVA, *P* < 0.05; 26.9% in leaders vs. 16.5% in followers). SIs are significantly higher in leaders than followers (two-tailed Mann-Whitney *U* test, *P* = 6.0 x 10^-171^). **c**, Probability correct of a support vector machine (SVM) decoder classifying choice trained on activity from 100 simultaneously recorded neurons in leaders or followers. Line and shading indicate mean ± SEM across animals (*n* = 4 leaders and 4 followers; 4-11 sessions per animal). Shuffled data combines animals from both groups. **d-g**, Calcium activity of four example neurons selective for trial-by-trial leadership dynamics. **d**,**e,** follower neurons; **f**,**g**, leader neurons. Left plots: each row is a trial, grouped by arrival order and sorted by arrival time. L, trials in which the recorded mouse leads; F, follows. Purple and green ticks denote leader and follower arrival, respectively. Right plots: mean ΔF/F ± SEM. Solid line, trials in which the recorded mouse leads; dashed line, follows. **h,i**, Selectivity indices for trial-by-trial leadership dynamics. Positive values indicate a preference for trials in which the recorded mouse arrives first. Each row is a significantly selective neuron from followers (**h**) or leaders (**i**) (permutation test, *P* < 0.05; see Methods). **j**, Proportions of reward zones chosen by each mouse when leading or following. Repeated measures ANOVA shows no significant effect of reward zone (*P* = 0.75; *n* = 8 animals), leadership role (*P* = 1.00), or their interaction (*P* = 0.30). **k**, Port choice does not differ between trials when mice lead or follow (two-tailed Wilcoxon signed-rank test, *P* = 0.95; *n* = 8 animals). **l**, Arrival speed of mice when leading or following. **m**, Arrival trajectories when leading or following. Insert shows plotting area (red square) relative to the full arena. **n**, Proportion of leadership-selective neurons modulated by reward expectation does not differ from overall population (two-tailed Wilcoxon signed-rank test, *P* = 0.64, *n* = 8 animals). **o**, Selectivity indices (SIs) for trial-by-trial leadership in follower (left) or leader (right) neurons. Shading indicates neurons with significant selectivity (permutation test, *P* < 0.05): 20.0% in followers (*n* = 6,995 neurons, 4 animals) and 14.5% in leaders (*n* = 8,395 neurons, 4 animals). Absolute SIs are significantly higher in followers than leaders (two-tailed Mann-Whitney *U* test, *P* = 6.3 x 10^-124^). A greater proportion of follower neurons prefer leading (12.8% vs. 7.2%; z-test for proportions, *z* = 10.9, *P* = 0), while more leader neurons prefer following (12.1% vs. 2.4%; *z* = 24.1, *P* = 0). **p**, SVM decoder classifying trial-by-trial leadership, trained on activity from 100 neurons in leaders or followers. Performance is measured as the area under the ROC curve (auROC). Lines and shading show mean ± SEM across animals (*n* = 4 leaders and 4 followers). Shuffled data pools both groups.

Notably, individual mPFC neurons tracked the recorded animal’s leadership status on a trial-by-trial basis, distinguishing whether the mouse arrived first or second (Fig. 4d-g). Some neurons responded selectively in trials when the recorded animal followed (Fig. 4d,f), while others prefer trials in which the animal led (Fig. 4e,g). These neurons were found in both followers and leaders. Interestingly, both groups showed a bias for the less frequent trial type, i.e., more neurons in followers prefer trials when they lead, whereas more leader neurons favor trials when following (Fig. 4h,i). To rule out confounds related to arrival order, we conducted several control analyses. First, reward zone and port choices did not differ between trials when mice arrived first or second, despite individual port preferences (Fig. 4j,k). Second, approach speed and trajectories to the reward zones were similar regardless of arrival order (Fig. 4l,m). Third, although animals that arrive first wait longer at the reward zone, only ∼20% of leadership-selective neurons exhibited ramping activity correlated with wait time, an indication of possible modulation by reward expectation, and this proportion did not differ from the overall population (Fig. 4n). After regressing out speed and position contributions from neural activity and stratifying trials by port choice, 20.0% of follower neurons and 14.5% of leader neurons remained selective for arrival order and more neurons in both roles continued to favor the less frequent trial types (Fig. 4o). A logistic regression decoder trained on mPFC activity from either leader or follower classified arrival order well above chance, with higher accuracy using follower activity (Fig. 4p).

These results demonstrate that mPFC neurons encode trial-by-trial leadership dynamics, with a higher proportion of selective neurons in followers than in leaders. Notably, both roles exhibited a biased representation favoring less frequent trial types: more follower neurons prefer trials in which the animals lead, while more leader neurons favor trials in which they follow. This pattern suggests that mPFC population activity not only encodes the momentary arrival order, but also reflects the animal’s enduring social role as a leader or a follower.

### An egocentric social value map of partner position in mPFC

Our behavioral analyses suggest that mice flexibly use both self and partner position to guide foraging decisions on a trial-by-trial basis. To examine the spatial coding, we analyzed mPFC activity in frames when both animals are outside the reward zones, a period when spatial information is most relevant and confounding reward signals are ruled out. In this context, spatial relationships can be represented in either allocentric (world-centered) or egocentric (self-centered) coordinates (Fig. 5a-d, top). We found that mPFC neurons predominantly encode the partner’s position in egocentric coordinates (examples in Fig. 5a-g, Supplementary Fig. 5a-c). The representative neuron in Fig. 5a-d (bottom) responded most strongly when the partner was located at the proximal front left of the recorded mouse, regardless of their absolute positions in the arena.

**Figure 5.**
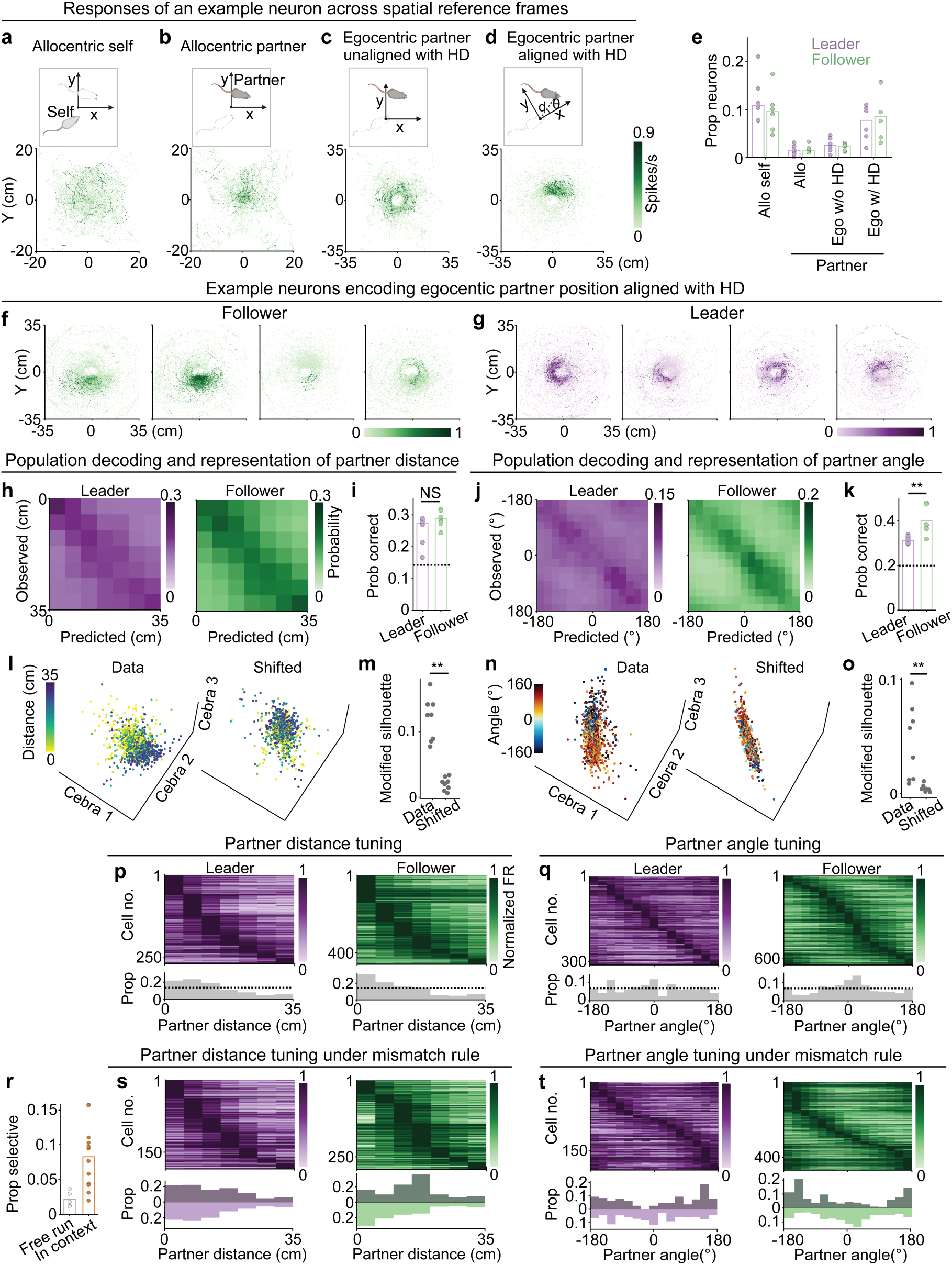
An egocentric social value map of partner position in mPFC. **a-d**, Top: Reference frames for representing self and partner positions. Allocentric coordinates of self (**a**) and partner (**b**) relative to the arena. Egocentric partner coordinates either unaligned (**c**) or aligned (**d**) with self heading; *d* and *θ* denote partner distance and angle, respectively. Bottom: An example mPFC neuron tuned to self or partner position in each reference frame. In the egocentric frames, the recorded mouse is positioned at the origin (**c,d**) and faces east (**d**). Activity is deconvolved spike rate from ΔF/F. Frames in which either mouse is within any reward zones are excluded. **e**, Proportion of neurons significantly tuned to each spatial representation. Each point represents one animal; bars indicate medians. Egocentric partner tuning aligned with heading does not differ between roles (two-tailed Mann-Whitney *U* test, *P* = 0.82, *n* = 6 animals per group), but is significantly greater than tuning for egocentric unaligned and allocentric partner representations (two-tailed Wilcoxon signed-rank test, both *P* = 4.9 x 10^-4^, *n* = 12 animals). **f,g**, Eight additional example neurons from followers (**f**) and leaders (**g**) showing egocentric encoding of partner position, format as in **d**. Activity represents the deconvolved spike rate from ΔF/F, normalized to each neuron’s maximum. **h**, Cross-validated confusion matrices for decoding partner distance from mPFC activity in a leader (left) or a follower (right), using a logistic decoder trained on 300 neurons per animal. **i**, Decoding accuracy for partner distance does not differ significantly between roles (two-tailed Mann-Whitney *U* test, *P* = 0.13; *n* = 6 animals per group). Each point represents one animal; bars indicate medians; dashed line denotes chance-level performance. **j**, Cross-validated confusion matrices for decoding partner angle from mPFC activity in a leader (left) or a follower (right). Format as in **h**. **k**, Decoding accuracy for partner angle is significantly higher using follower mPFC activity (two-tailed Mann-Whitney *U* test, *P* = 0.0087; *n* = 6 animals per group). Format as in **i**. **l**, CEBRA multi-session embeddings for a representative mouse using data (left) or time-shifted (right) partner distance labels. **m**, Modified silhouette scores comparing embeddings from data versus shifted partner distance (one-tailed Wilcoxon signed-rank test, *P* = 0.0039; *n* = 8 animals). **n**, CEBRA embeddings for partner egocentric angle. Format as in **l**. **o**, Modified silhouette scores comparing embeddings from data versus shifted partner egocentric angle (*P* = 0.0039; *n* = 8 animals). **p**, Tuning curves of partner distance-selective neurons in leaders and followers. Heatmaps show normalized activity of individual neurons sorted by preferred distance; histograms below indicate the preferred bins. Dashed lines indicate the chance level (1/7). Both groups exhibit significant overrepresentation of neurons tuned to near distances (126 of 268 neurons in the two proximal bins in leaders, binomial test against 2/7, *P* = 5.7 × 10^-11^; 250 of 480 in followers, *P* = 1.0 × 10^-27^). **q**, Tuning curves of egocentric partner angle-selective neurons, as in **p**. Followers show a prominent central peak (215 of 644 neurons in the three center bins, binomial test against 3/15, *P* = 1.2 × 10^-15^), whereas leaders do not (70 of 311 neurons, *P* = 0.15). **r**, Proportion of neurons selective for egocentric partner position is significantly lower in animals freely interacting outside the task context (two-tailed Mann-Whitney *U* test, *P* = 0.0023, *n* = 5 free-run animals, 12 task-engaged animals). **s**, Tuning curves for partner distance in pairs retrained under the mismatch rule. Mirrored histograms show tuning under the original match rule. Leader tuning remains unchanged (two-tailed Kolmogorov-Smirnov test, *P* = 0.52), whereas follower tuning shifts significantly toward longer distances (*P* = 1.6 × 10^-11^). **t**, Tuning curves for egocentric partner angle after retraining under the mismatch rule. Both roles show significant shifts from frontal to rear angles (*P* = 8.8 × 10^-4^ and 8.8 × 10^-17^, respectively).

We identified mPFC neurons selective for self or partner position in either allocentric or egocentric coordinates using three criteria: spatial information content, spatial coherence, and within-session stability (Methods). A subset of neurons (10.9%) represented the animal’s own position in allocentric coordinates. Notably, 8.3% of neurons encoded the partner’s position in an egocentric reference frame—approximately four times more than those tuned to partner position in allocentric coordinates or egocentric coordinates not aligned with the animal’s own heading (Fig. 5e). These proportions were similar between leaders and followers, indicating that spatial encoding of self and partner is preserved across roles. At the population level, logistic decoders trained on mPFC activity predicted both the partner’s distance and egocentric angle, corresponding to the radial and angular components of partner position in a polar coordinate system centered on the recorded mouse (Fig. 5h-k, Supplementary Fig. 6a,b). Decoding accuracy for partner distance did not differ between roles (Fig. 5i), but decoding of partner angle was significantly more accurate in followers, consistent with their greater need to orient toward leaders (Fig. 5k).

We then used CEBRA^47^, a contrastive learning framework that extracts low-dimensional structure in neural activity by isolating variance specifically related to the behavioral variables of interest. This approach allowed us to examine how partner distance and egocentric angle are organized in mPFC population activity by generating latent embeddings that capture these task-relevant features. Using supervised CEBRA multi-session models trained with continuous labels for partner distance or angle, we projected held-out neural activity into a 3D latent space. These embeddings revealed structured organization for both partner distance and angle (Fig. 5l,n). Compared to control embeddings trained with time-shifted labels, embeddings trained with true labels produced significantly better structure across animals (Fig. 5m,o). These findings corroborate the decoding results and suggest that partner information is organized into a continuous, behaviorally relevant space in mPFC for cooperative foraging.

We next examined how these egocentric partner-selective neurons are tuned to the distance and angle of the other mouse. In both leaders and followers, mPFC neurons selective for partner distance showed a strong preference for close-range positions (Fig. 5p), reflecting the behavioral strategy of maintaining proximity during cooperation. Remarkably, while the tuning profile for egocentric partner angle was relatively uniform in leaders, follower neurons showed a pronounced bias toward angles near zero degrees (Fig. 5q)—where the partner is directly in front—consistent with the follower’s need to orient toward and follow the leader.

The asymmetric and skewed tuning suggests that mPFC neurons do not simply form a static spatial map of partner position. Rather, their activity may reflect the behavioral relevance or value associated with specific partner positions. To test this, we conducted two experiments. First, we imaged mPFC activity in mice freely interacting in the same environment after water satiation, outside the task context. Under these conditions, the proportion of egocentric partner-selective neurons was significantly reduced (Fig. 5r). Second, we retrained a subset of well-trained pairs on a reversed “mismatch” rule, in which mice were rewarded only when they entered different zones (Methods). Mice successfully adapted and performed the new task with high accuracy (Supplementary Fig. 6c). In this new context, mPFC neurons exhibited markedly different tuning. Preferred partner distance in followers shifted toward longer ranges (Fig. 5s), while partner angle tuning decreased around 0° and increased near ±180° in both roles (Fig. 5t). These changes are consistent with a strategy of avoiding close frontal proximity, as required by the mismatch rule. Together, these findings show that mPFC neurons encode partner position in an egocentric reference frame, with follower neurons displaying stronger and more directional tuning. These representations are not static spatial maps but flexibly reflect the behavioral value of partner positions.

### Model-inferred social value aligns with role-specific mPFC representations

We developed a novel multi-agent inverse reinforcement learning (MAIRL) ^48^ algorithm, to further examine animals’ latent value functions, or internal representations of goals and preferences guiding cooperative behavior (Methods). MAIRL decomposes the joint value function into marginal and interaction maps, addressing the exponential increase in computational complexity that arises with multiple agents. Based on observed animal trajectories, we infer the best-fitting social value functions, comprising separate components for behavioral variables such as self and partner allocentric position, partner distance, and egocentric partner angle (Fig. 6a). During inference, we determine which components to include based on unbiased model selection, quantified by the log-likelihood gain relative to a base model.

**Figure 6.**
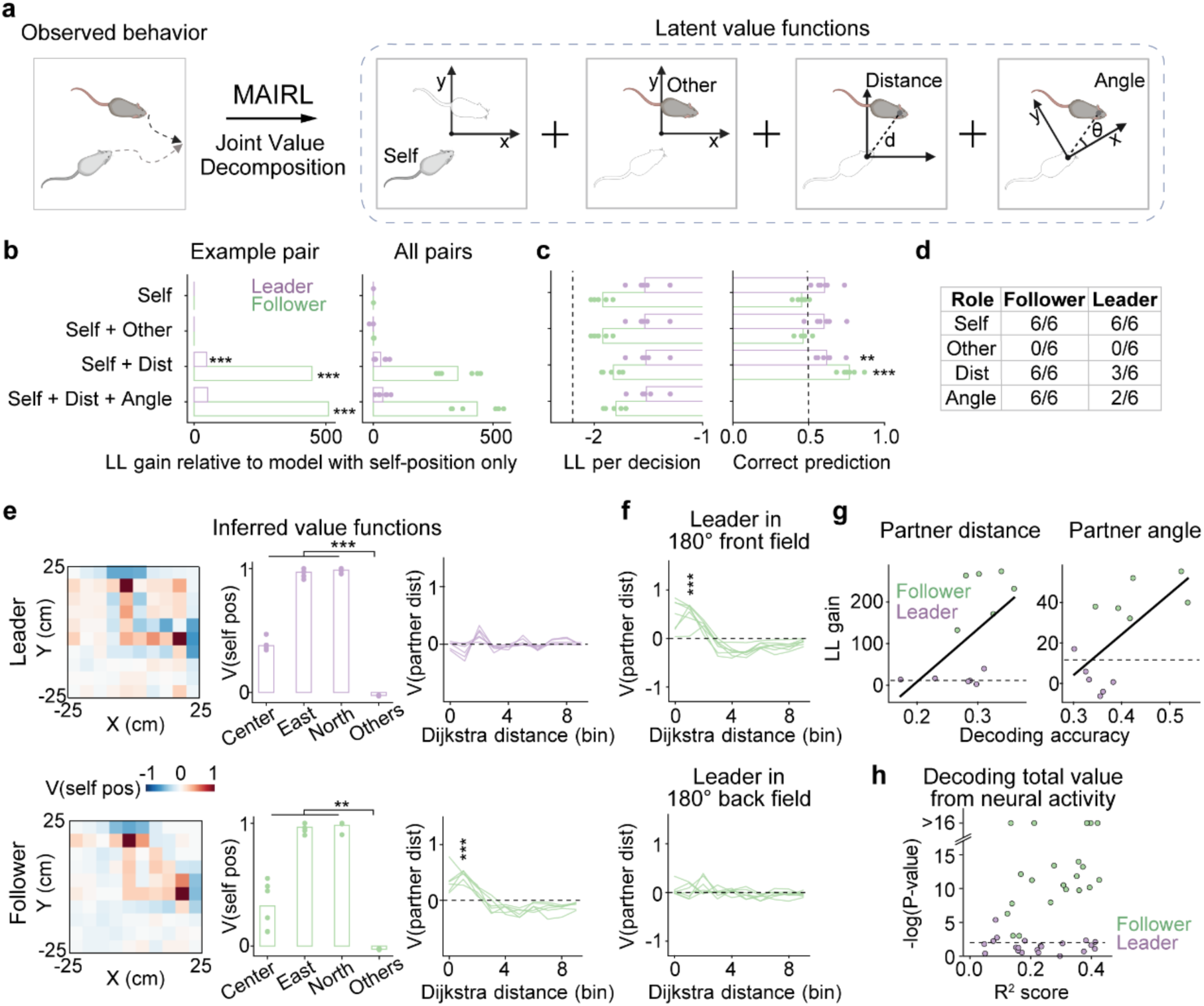
MAIRL-inferred social value aligns with role-specific mPFC representations. **a,** Schematic illustrating inference of latent value functions from observed animal trajectories using MAIRL. The composite value function is modeled as a sum of components corresponding to allocentric self-position, allocentric partner-position, self-partner distance, and egocentric partner angle. Components are included in the final model based on hypothesis testing and model selection, retaining only those that significantly improve model fit. **b,** Model fit improvement for an example pair (left) and across all pairs (right), measured as log-likelihood (LL) gain relative to a baseline model including only self-position. Adding allocentric partner position did not improve model fit for either role. In contrast, including partner distance significantly improved model performance in both leader and follower (chi-square test of nested models, same below; \*\*\**P* = 2.5 x 10^-16^ and 6.5 x 10^-186^, respectively), with a further improvement in follower when egocentric partner angle was added (\*\*\**P* = 4.2 x 10^-22^). **c**, Behavioral prediction from simulated trajectories using inferred value functions. Left: Log-likelihood (LL, in nats per decision) across all leader–follower pairs for different models. The dashed line indicates the LL of a baseline model assuming equal action probabilities. Right: Accuracy for predicting the reward zone chosen by the animals from model-simulated trajectories. The “self + distance + angle” model could not be simulated because angle was only available as an input for the model. For both leader and follower animals, the “self + distance” model significantly outperforms the baseline model in predicting reward zone choices (one-sample *t*-test compared to baseline prediction value, \*\**P* = 0.0064 for leaders and \*\*\**P* = 1.6 x 10^-4^ for followers). **d,** Number of animals (out of total) showing a significant improvement in model fit when each component was sequentially added. The model with self-position was compared to a constant model that assumes equal probability of all actions, and subsequent components were tested against the best-fitting simpler model. **e**, Inferred value functions in correct trials for leaders (top row) and followers (bottom row). Left, example self-position value map in allocentric space from one animal. Middle, value assigned to the initiation spot and to the east and north reward ports, and the mean value of all other locations across animals. Both leaders and followers assign significantly higher value to initiation and reward locations (paired *t*-test, all \*\*\**P* < 4.2 x 10^-6^ for leaders, *n* = 6 animals; all \*\**P* < 0.0074 for followers). Right, value functions for partner distance across animals. Followers assign significantly higher value to proximal partner locations than leaders (*t*-test comparing values at distance = bin 1, *P* = 7.7 x 10^-6^; *n* = 6 animals per group). **f**, Inferred value maps for partner distance in followers, conditioned on the leader’s egocentric angle. Value peaks at close range when the leader is in front (left), but diminishes when behind (right) (paired *t*-test, *P* = 3.3 x 10^-4^; *n* = 6 animals). **g,** Correlation between model fitness gain from including partner distance (left) or angle (right) and the decoding accuracy for each term using mPFC activity (Pearson’s correlation, *n* = 12; distance: *R* = 0.54, ^#^*P* = 0.069; angle: *R* = 0.67, \**P* = 0.018). Dashed lines indicate the threshold for statistical significance (*P* < 0.01) in fitness gain. **h,** Coefficient of determination (R^2^ score) for predicting the inferred composite value (sum of all value function components) from mPFC population activity, with corresponding P-values shown on the y-axis in log scale. Each point represents one session (*n* = 21 leader, 22 follower sessions). P-values were computed by comparing true R^2^ scores to null distributions generated by circularly shifting the value labels (Methods). Dashed line indicates the threshold for statistical significance (*P* < 0.01). Predictions based on follower activity consistently outperform those based on leader activity.

Both leaders and followers primarily valued their own position and partner distance, with followers exhibiting significantly greater log-likelihood gain for the latter—consistent with their need to maintain proximity during coordination (Fig. 6b). Neither role placed value on the partner’s allocentric position. Notably, followers showed further improvements in model fit when egocentric partner angle was included, whereas leaders exhibited less sensitivity. To evaluate model fitting, we simulated foraging trajectories using the inferred value functions and predicted the reward zone choice by the animals. Including partner distance in the model significantly improved the predictions (Fig. 6c). For all followers, the best-fitting models required both partner distance and egocentric angle, whereas only a subset of leaders showed such dependencies (Fig. 6d).

We next examined the composite value functions, which consist of separate value maps for the animal’s own allocentric position and the distance to its partner (Fig. 6e). The value maps of self-position displayed hotspots at the initiation spot and reward ports, consistent with their task relevance (Fig. 6e, left and middle). In contrast, the value maps of partner distance revealed a prominent peak in followers, but not in leaders, when the partner was nearby (Fig. 6e, right). This peak was significantly stronger when the leader was positioned in front than behind (Fig. 6f), reflecting a key follower strategy of maintaining close, front-facing proximity to the leader. These value representations were diminished during error trials (Supplementary Fig. 7a) and under the “mismatch” rule (Supplementary Fig. 7b).

We next asked whether the inferred value functions are reflected in the mPFC population activity. Notably, the improvement in model fit from including partner angle was correlated with the decoding accuracy for partner angle from neural activity across animals, with a similar trend observed for partner distance (Fig. 6g). To further test this link, we attempted to decode the inferred value functions directly from mPFC activity. Decoding was consistently above chance in all follower recordings, while only a subset of leader sessions showed significant decoding (Fig. 6h, Supplementary Fig. 7c), suggesting that value representations are more robustly encoded in follower mPFC. These results validate our inference approach and establish a quantitative correspondence between latent value functions and neural representations.

Together, these findings show that MAIRL uncovers the latent structure of role-based social decision-making. Differences in value functions between leader and follower—especially the greater value followers assign to close, front-facing proximity to partners and their sensitivity to task context—closely mirror the mPFC activity. Crucially, the ability to decode these value functions from mPFC supports the idea that this region encodes a dynamic, role-specific social value map uncovered by MAIRL.

## Discussion

While studies in humans, non-human primates, and other species have begun to explore the neural basis of cooperative behavior^28,46,49–52^, a robust and ethologically grounded paradigm remains lacking in mice. We developed a novel behavioral paradigm that combines spatial coordination with free social interaction, addressing this gap and overcoming key limitations of previous approaches^46,53^. Our findings demonstrate that mice not only perform this task robustly but also develop emergent social roles and exhibit flexible and reciprocal coordination. These roles are stable across sessions, and the timing and strength of this emergent leadership dynamic strongly predicts learning. The social roles exhibit asymmetric yet reciprocal influence: leaders exert a greater impact on followers’ decisions, but followers also shape the behavior of leaders. Interestingly, although initiators are often positioned farther from the nearest active zone than responders, we did not observe a consistent division of labor such that initiators are always followers. This suggests that while physical proximity may shape moment-by-moment decisions, the role differentiation may be governed by more stable traits such as motivation.

Importantly, leadership and followership roles are not associated with dominance hierarchy, indicating that they reflect a distinct form of social organization shaped by cooperative demands rather than preexisting power dynamics. Our approach aligns well with rodents’ natural foraging behaviors, highlighting the importance of studying freely interacting group behavior^28,54^ and suggesting that tasks emphasizing spatial coordination may better exploit the social cognitive capacities of mice. Forward reinforcement learning modeling using MARL revealed that behavioral asymmetries and social roles such as initiator and leader can emerge spontaneously among initially identical agents, likely driven by small stochastic differences in learning. This convergence between emergent roles in simulation and mice underscores the capacity of shared incentives and coordination demands to give rise to social roles.

Multiple lines of evidence confirm that our cooperative foraging task captures genuine mouse social behavior. First, the strong reciprocity between leader and follower roles, as described above, reflects dynamic mutual influence rather than unilateral control. Second, we observe significant differences in learning between males and females, suggesting that the task is sensitive to innate, sex-dependent factors. While such differences are not definitive on their own, they are consistent the well-documented influence of sex on many forms of social behavior. Third, when well-trained animals are separated and paired with new partners, they tend to preserve their original roles, yet require time to adapt to the unfamiliar partner. This indicates that leader and follower roles are stable social identities that persist across scenarios but can also adapt to individual idiosyncrasies. Fourth, well-trained animals exhibit a rich repertoire of behavioral motifs, the composition and frequency of which vary across pairs and roles, revealing both behavioral flexibility and social individuality. Finally, neural recordings show that mPFC in both animals encodes trial-by-trial leadership dynamics and egocentric position of the partner. This suggests that both roles are actively engaged in the task and maintain internal models of their partner and their own, reinforcing the fundamentally social nature of this interaction.

While primate studies have implicated the frontal cortex in social cognitive processes such as shared experience and opponent prediction, our findings provide novel evidence that cooperative behavior in rodents critically depends on the mPFC in a reciprocal and role-specific manner. Both long-lasting chemogenetic and transient optogenetic inactivation of the mPFC in leaders and followers reduced cooperation rate, underscoring its essential role in supporting cooperative behavior. Notably, when the mPFC was inactivated in only one animal—regardless of role—the partner often compensated through adaptive adjustments, highlighting reciprocal flexibility as a core feature of cooperative behavior. This adaptability may explain why transient optogenetic silencing of a single animal did not impair performance. Interestingly, while leader-follower roles were disrupted by chemogenetic inactivation, they remained intact under optogenetic silencing. The former likely interferes with the maintenance of social roles through sustained disruption of mPFC function, whereas the latter may be insufficient to destabilize the underlying organizational structure, as these roles are embedded in the animals’ learned strategies.

The discovery that mPFC encodes trial-by-trial leader-follower dynamics reveals a novel dimension of social representation. While primate studies have implicated the frontal cortex in processing self- and other-related information, they have not addressed emergent social roles that structure group behavior. In rodents, prior work has primarily focused on innate roles such as dominance hierarchy. By contrast, our findings demonstrate that mPFC neurons dynamically encode whether an animal is leading or following on a trial-by-trial basis. This suggests that mPFC supports real-time tracking of self-other relationships, potentially enabling credit assignment^55^ and facilitating the learning and consolidation of social roles. Furthermore, the observation that neurons preferentially encode less frequent leading or following states within each role indicates that mPFC activity reflects the animal’s enduring social role identity. This may reflect a neural encoding of social prediction error or deviations from expected norms, mechanisms that may be critical for adapting behavior in dynamic social contexts.

mPFC recordings revealed a dynamic social value map that encodes the partner’s position in an egocentric frame of reference. Prior work in the hippocampus has identified “social place cells” that represent conspecific positions in allocentric coordinates during observational tasks^56,57^. A more recent study reported multiplexed hippocampal encoding of partner location using both allocentric and egocentric frames^58^. The prelimbic cortex was also shown to encode a combination of social and allocentric spatial information^59^. By contrast, our data show a fourfold enrichment in mPFC neurons tuned to egocentric partner position relative to other spatial coordinates. Population-level analysis further demonstrated that partner position is organized into continuous, behaviorally relevant latent spaces. Critically, this spatial coding in mPFC is not static. Rather, it is role-specific, value-based, and dynamically modulated by task demands. mPFC neurons in followers show enhanced tuning to close, front-facing proximity, which adapts flexibly with changes in task rules, supporting its role as a dynamic social value map rather than a static spatial representation. This positions the mPFC as a key node in computing social value by integrating egocentric partner information with the animal’s own role and behavioral strategy.

Our inverse reinforcement learning modeling demonstrates the power of MAIRL to disentangle the latent social and non-social components of value representations guiding social decision-making. By decomposing value functions into interpretable components including self-position and partner distance and angle, MAIRL enables a principled dissection of the distinct computations involved in cooperative behavior. Importantly, the value functions inferred from observed behavior generate testable hypotheses about how the brain assigns value to distinct elements of social interaction. We find that these inferred value functions are not only dynamic and role-dependent, but also qualitatively recapitulate and are quantitatively decodable from mPFC population activity. This corroborates both the validity of our model and the encoding of social value in mPFC, establishing a strong link between computational inference and neural representation. Further, MAIRL provides a flexible and generalizable framework that can be extended to more agents and other types of social interactions, offering a path to study complex group behavior including the ones involving social inference and recursive reasoning^60^. This approach thus lays the groundwork for formalizing and probing latent cognitive variables that shape collective behavior across species.

Our paradigm establishes a critical foundation for mechanistic studies of cooperation in mice, offering an ethologically grounded and experimentally tractable model that supports the full range of modern circuit-level tools. In parallel, our MAIRL algorithm introduces a novel computational framework that infers latent value functions from observed behavior, providing a principled, interpretable decomposition of social and non-social decision variables. Together, this integrated experimental and modeling approach allows for direct mapping between behavioral strategies, latent cognitive computations, and neural representations. It opens new avenues for probing core questions in social cognition such as norm adherence, role differentiation, and social inference and for testing the “social brain” hypothesis^61–63^. Our approach can synergize with ecological and psychological studies of group behavior by offering neural and algorithmic insights into the emergence and function of social roles. Beyond mice, this framework can be adapted to study more complex group behavior, including multi-agent interactions and strategic games in other animal models and human research. Our findings may offer insights into social dysfunction in neuropsychiatric disorders and inform biologically inspired approaches to adaptive multi-agent AI systems.

---

**Supplementary Figure 1.**
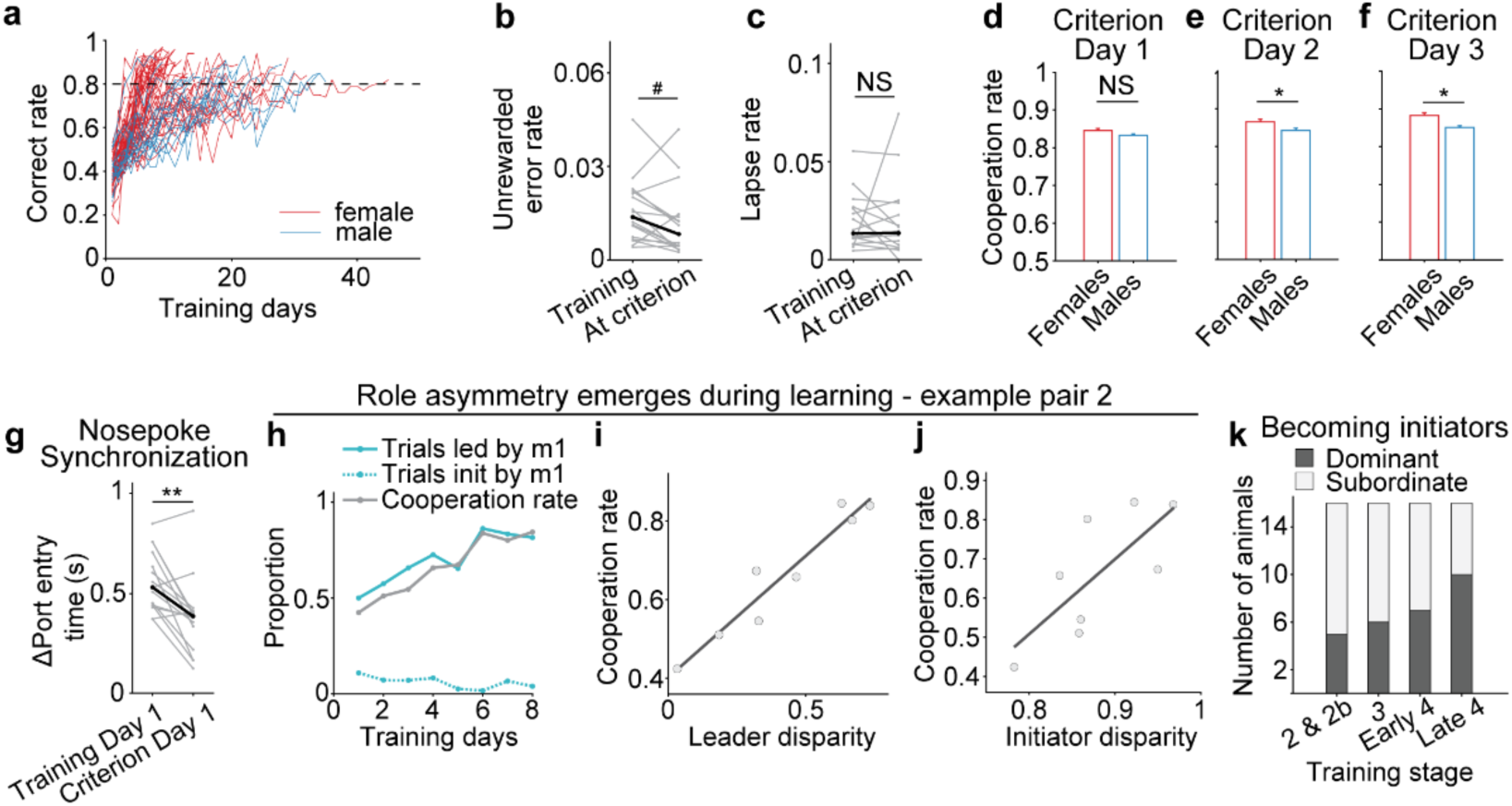
Training progression and emergence of social roles in cooperative foraging. **a**, Learning curves for female (red, *n* = 55 pairs) and male (blue, *n* = 21 pairs) pairs. The dashed line indicates the 80% correct training criterion. **b**, Unrewarded error rates during the learning (median = 0.014, IQR = 0.015) and well-trained (median = 0.0083, IQR = 0.0098) stages. Error rates show a decreasing trend (Wilcoxon signed-rank test, ^#^*P* = 0.088, *n* = 15 pairs). **c**, Lapse rates during the learning (median = 0.014, IQR = 0.017) and well-trained (median = 0.014, IQR = 0.019) stages did not significantly differ (Wilcoxon signed-rank test, *P* = 0.50, *n* = 15 pairs). **d-f**, Correct rates for female and male mouse pairs in the first (**d**), second (**e**), and third (**f**) sessions at the training criterion. Females exhibit better performance than males. two-tailed Mann-Whitney *U* test: *P* = 0.19 (**d**), \**P* = 0.028 (**e**), \**P* = 0.016 (**f**). Data are mean ± SEM. **g**, Mean time difference in nose poke at the reward port between partners, measured in correct trials comparing the first training day to the first criterion day. Black line indicates the median (Wilcoxon signed-rank test, \*\**P* = 0.0030; *n* = 15 pairs). **h-j,** Role differentiation and learning in a second example pair with faster learning than the pair shown in Fig.1 d-f. **h,** Cooperation rate alongside the proportion of trials led (solid lines) or initiated (dashed lines) by one mouse across training days. **i,** Correlation between leader disparity and cooperation rate. Pearson’s correlation: *R* = 0.96, *P* = 1.3×10^-4^, 8 sessions. **j,** Correlation between initiator disparity and cooperation rate. *R* = 0.74, *P* = 0.034. **k**, Number of dominant and subordinate mice identified as initiators across training stages. Proportions are not significantly different from chance (two-tailed binomial test against 0.5, *P* > 0.21 for all four stages). No significant differences in proportions across stages (χ² (3, *n* = 64) = 3.56, *P* = 0.31).

**Supplementary Figure 2.**
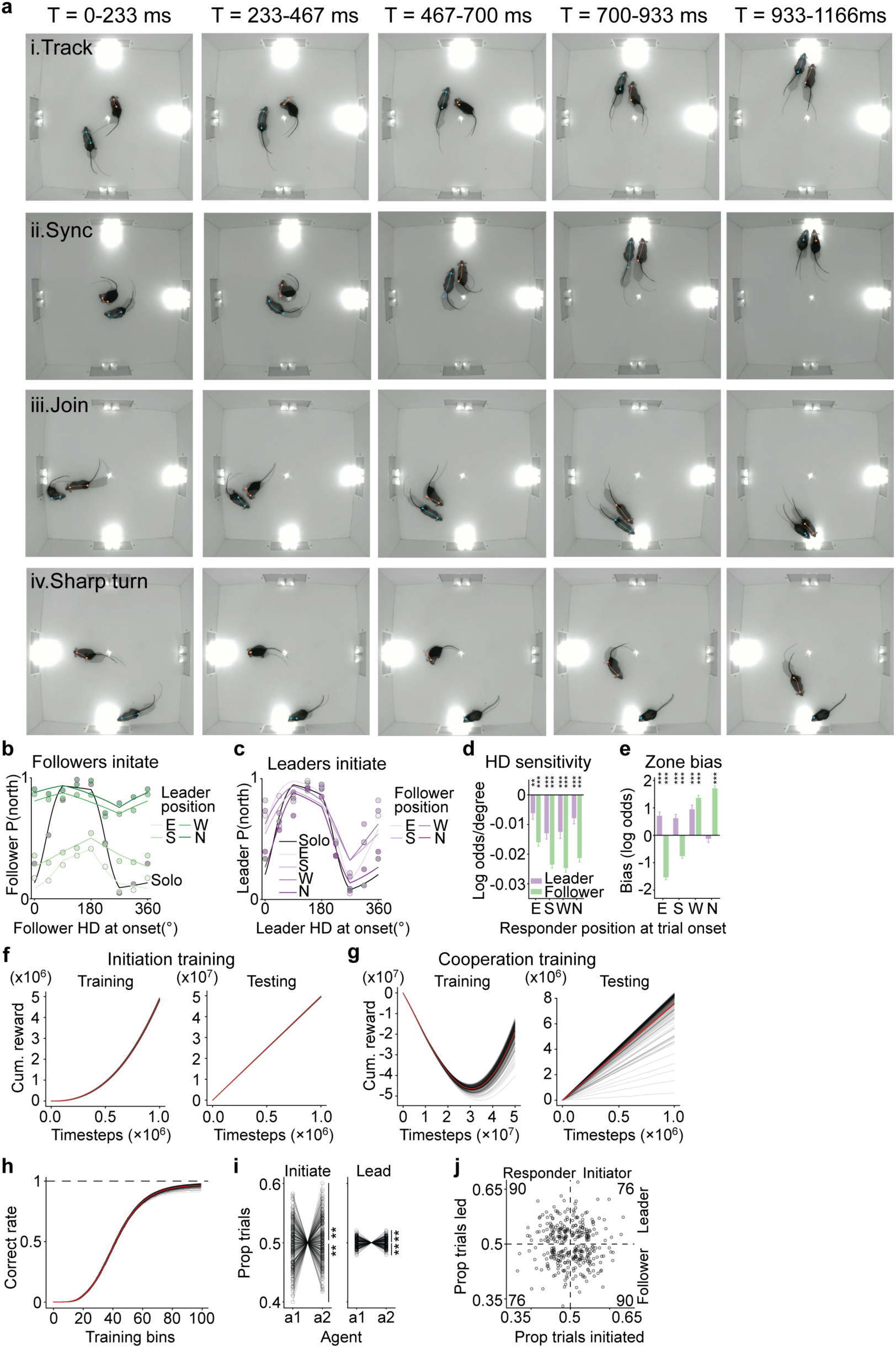
Social influence on decision-making in mice and learning dynamics in simulation. **a**, Time-binned example video frames (233 ms intervals) of the behavioral motifs. **b**, Follower choice plotted against its heading at trial onset. Each point indicates the proportion of trials in which the follower selected north, binned in 45° intervals centered on the x-axis values. Curves show logistic fits of P(north) as a function of follower heading, conditioned on leader’s position at trial onset (in the east, south, west, or north), or during solo foraging sessions. **c**, Leader choice as a function of its heading at trial onset; format as in **b**. **d,e**, Beta estimates from logistic regression showing how responder position affects initiator’s sensitivity to its own heading (**d**) and initiator’s bias in reward zone choice (**e**); positive values indicate bias toward north, negative toward east. **d**, When leader is the responder, its presence significantly reduces follower’s sensitivity (all \*\*\**P* < 1.6 × 10^-35^; *n* = 8 pairs, 4,517–4,894 trials) and vice versa (east: \*\**P* = 0.0028; south, west, north: \*\*\**P* < 6.7 × 10^-5^; *n* = 8 pairs, 3,536–3,624 trials) **e**, Leader in the east and south decreases follower bias, while in west and north increases it (all \*\*\**P* < 1.5 × 10^-20^). Follower in the east, south and west significantly increases leader bias (all *\*\*\*P* < 1.1 × 10^-5^). **f,** Cumulative reward over 1 x 10^6^ timesteps during initiation training and testing, where both agents are rewarded whenever either enters the central initiation zone. In training (left), extensive early exploration transitions to exploitation, producing a convex-upward increase in cumulative reward. Testing (right) shows linear increase under exploitation only. Grey lines indicate individual pairs; red lines indicate the mean with a thin band of SEM. **g,** Similar trend in cooperation training, where agents are rewarded only when arriving at the same target zone. However, early mismatches drive cumulative reward below zero in training (left), resulting in a U-shaped recovery as coordination is learned over more than 5 x 10^7^ timesteps. Grey lines indicate individual pairs; red lines indicate the mean with a thin band of SEM. **h**, Correct rate across cooperative training. Thin grey lines show each pair’s proportion of correct trials binned into 100 evenly spaced blocks, revealing a sigmoidal rise from near-zero to ceiling performance. **i,** Role-adoption asymmetry during cooperative training. Circles with connecting lines show each pair’s proportions of trials initiated and led by agent 1 versus agent 2 for pairs with significant bias (binomial test, *P* < 0.01); grey lines without circles near the center indicate pairs without significant bias. **j**, Proportion of trials initiated (x axis) and led (y axis) by individual agents during testing. Only agents with statistically significant bias in both initiation and leadership are shown and tallied by quadrant. The proportion of leaders who are also initiators does not differ significantly from chance (binomial test against 0.5, *P* = 0.31).

**Supplementary Figure 3.**
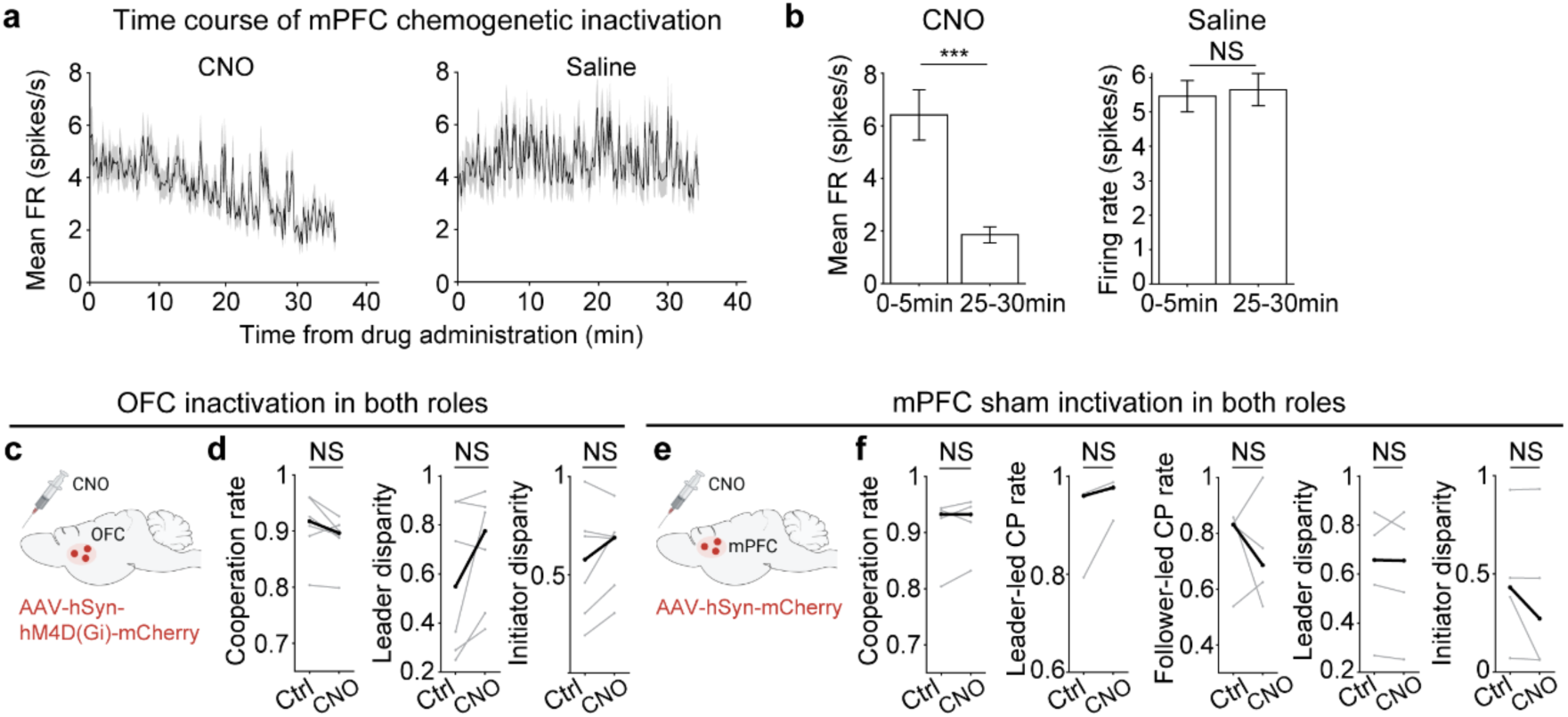
Validation and specificity of mPFC inactivation. **a**, mPFC population firing rates from example recording sessions following CNO (left, *n* = 49 units) or saline (right, *n* = 62 units) injection. Black line shows the mean; grey shading indicates the SEM. **b**, mPFC mean firing rates during the 0–5 min and 25–30 min time windows after CNO (left) or saline (right) injections. Neural activity significantly decreased after CNO (mean ± SEM; paired t-test, \*\*\**P* = 8.3 × 10^-6^; *n* = 2 animals, 82 neurons), but not after saline injection (*P* = 0.30; *n* = 2 animals, 190 neurons). **c**,**d**, Bilateral OFC inactivation in both leaders and followers did not affect cooperation rate (two-tailed Wilcoxon signed-rank test, *P* = 0.16, *n* = 6 pairs), leader disparity (*P* = 0.16), or initiator disparity (*P* = 0.44). **e**,**f**, Bilateral mPFC sham inactivation in both leaders and followers did not affect cooperation rate (two-tailed Wilcoxon signed-rank test, *P* = 0.62, *n* = 4 pairs), leader-led cooperation rate (*P* = 0.12), follower-led cooperation rate (*P* = 1.00), leader disparity (*P* = 0.88), or initiator disparity (*P* = 0.38).

**Supplementary Figure 4.**
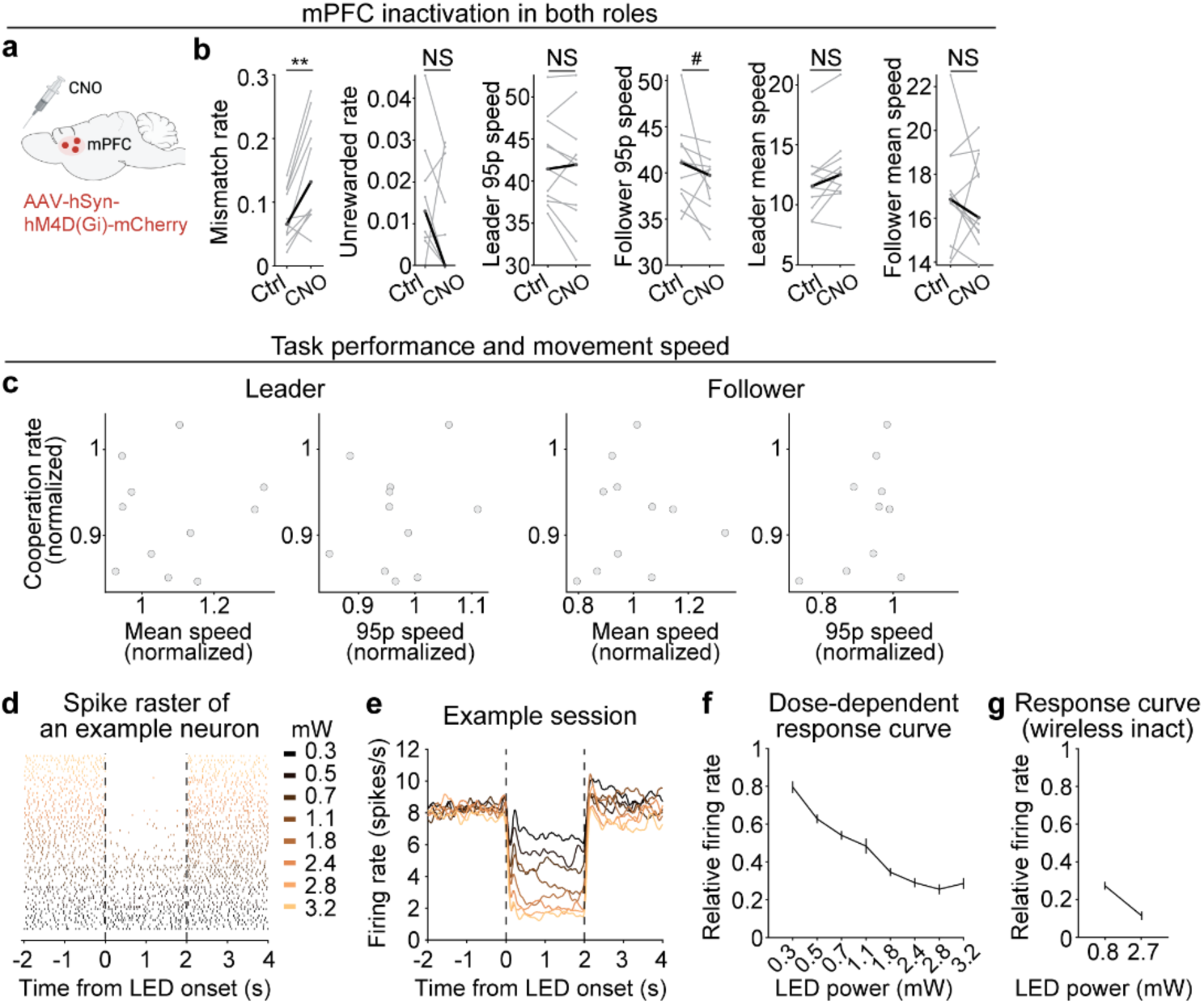
Behavioral specificity and optogenetic validation of mPFC inactivation. **a**,**b**, mPFC inactivation in both leaders and followers increased the mismatch rate (Wilcoxon signed-rank test, \*\**P* = 0.0029, *n* = 11 pairs), but did not affect unrewarded error rate (*P* = 0.19, *n* = 11 pairs), leader 95th percentile speed (*P* = 0.24), follower 95th percentile speed (^#^*P* = 0.083), leader mean speed (*P* = 0.10) or follower mean speed (*P* = 0.76); all speeds are in cm/s. **c**, Lack of correlation between cooperation rate and movement speed of leaders and followers, with both variables normalized to the control session the day before the inactivation session. No significant correlations were found for mean speed (leaders: Pearson’s correlation, *R* = 0.060, *P* = 0.86, *n* = 11 pairs; followers: *R* = 0.090, *P* = 0.79) or 95th percentile speed (leaders: *R* = 0.18, *P* = 0.59; followers: *R* = 0.25, *P* = 0.46). **d**, Spike raster of a representative mPFC neuron expressing KALI-1 under 590 nm LED illumination across power intensities (0.3–3.2 mW). **e**, mPFC mean firing rate across light intensities from an example recording session (*n* = 66 units). **f**, Normalized mPFC mean firing rate under LED illumination across power intensities (mean ± SEM; *n* = 135 units from 2 animals). **g**, Normalized mPFC mean firing rate under wireless LED illumination (mean ± SEM; *n* = 44 units from 1 animal).

**Supplementary Figure 5.**
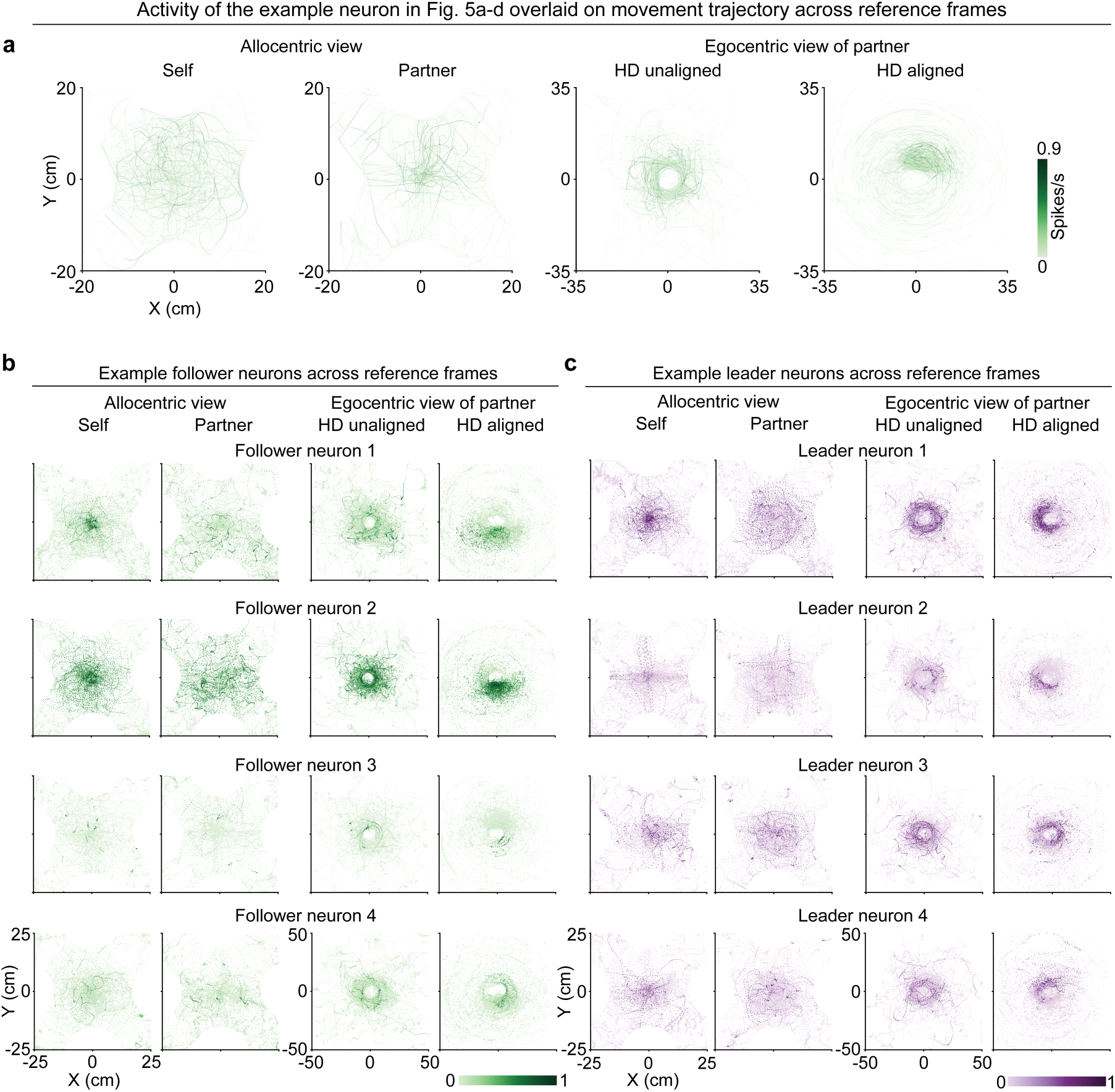
Spatial tuning of example neurons across allocentric and egocentric reference frames. **a**, Neural activity of the same example neuron shown in Fig. 5a–d, overlaid on movement trajectories across spatial reference frames: allocentric view of self and partner, and egocentric view of the partner (either unaligned or aligned with self heading). **b**, Example follower neurons showing spatial tuning across the reference frames. Format as in Fig. 5a-d. **c**, Example leader neurons showing spatial tuning across the reference frames. Format as in **b**.

**Supplementary Figure 6.**
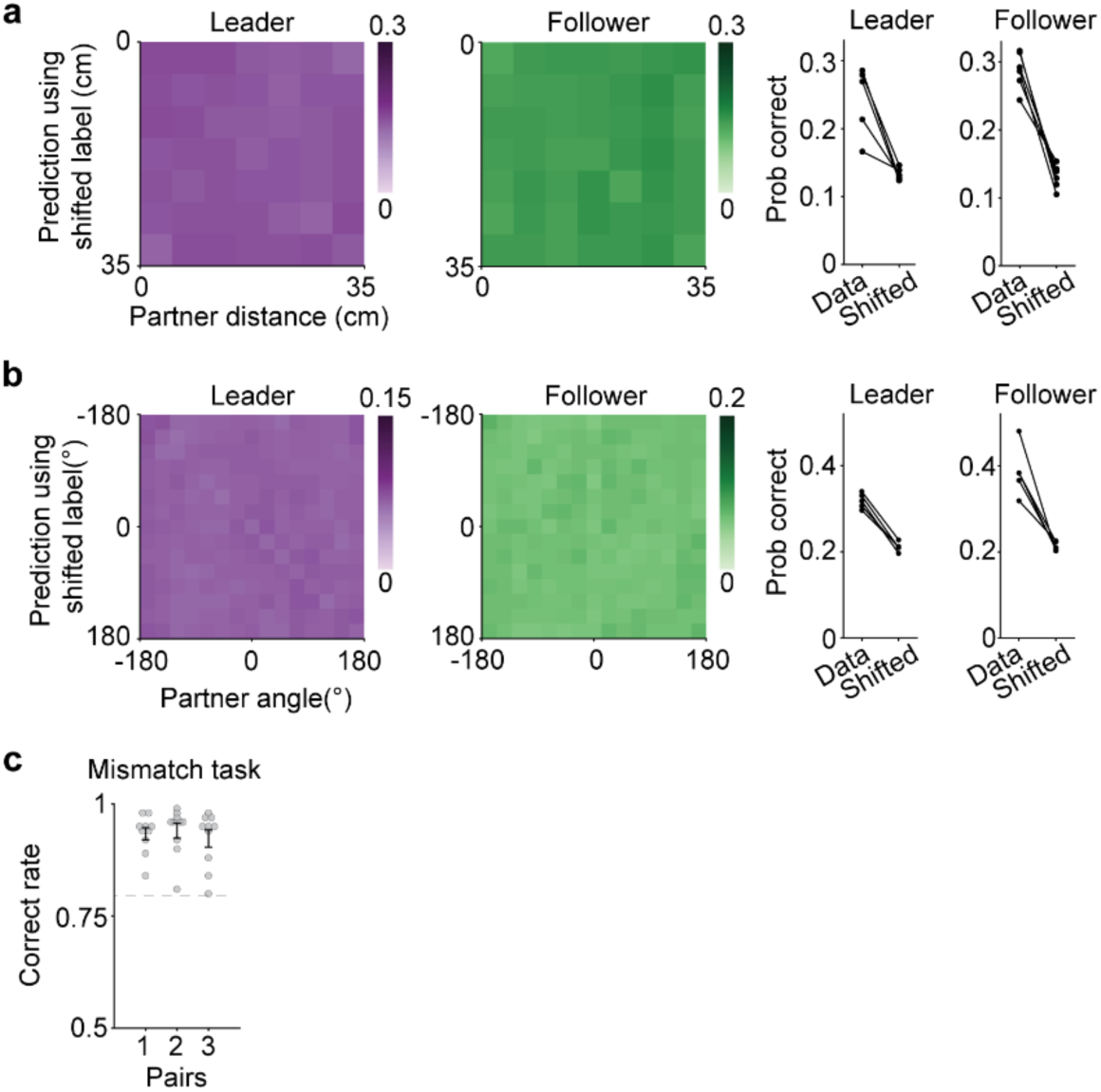
Decoding validation and behavioral performance under the mismatch rule. **a**, Control decoding accuracy of partner distance using circularly time-shifted labels. Left plots: Confusion matrices show the probabilities of correct predictions for leaders and followers. Right plots: Decoders trained with true labels significantly outperform those trained with shifted labels (Mann–Whitney U test; leaders, *P* = 0.011; followers, *P* = 2.1 × 10^-4^; *n* = 6 animals per group). Each point represents one animal. **b**, Control decoding accuracy of egocentric partner angle using circularly time-shifted labels. Format as in **a**. Decoders trained with true labels significantly outperformed the shifted labels (Mann–Whitney U test; leaders, *P* = 1.2 × 10^-4^; followers, *P* = 0.0045; *n* = 6 animals per group). **c**, Behavioral performance under rule reversal (reward on mismatch; mean ± SEM; *n* = 3 pairs, 10 sessions each). Dashed line indicates the 80% correct training criterion.

**Supplementary Figure 7.**
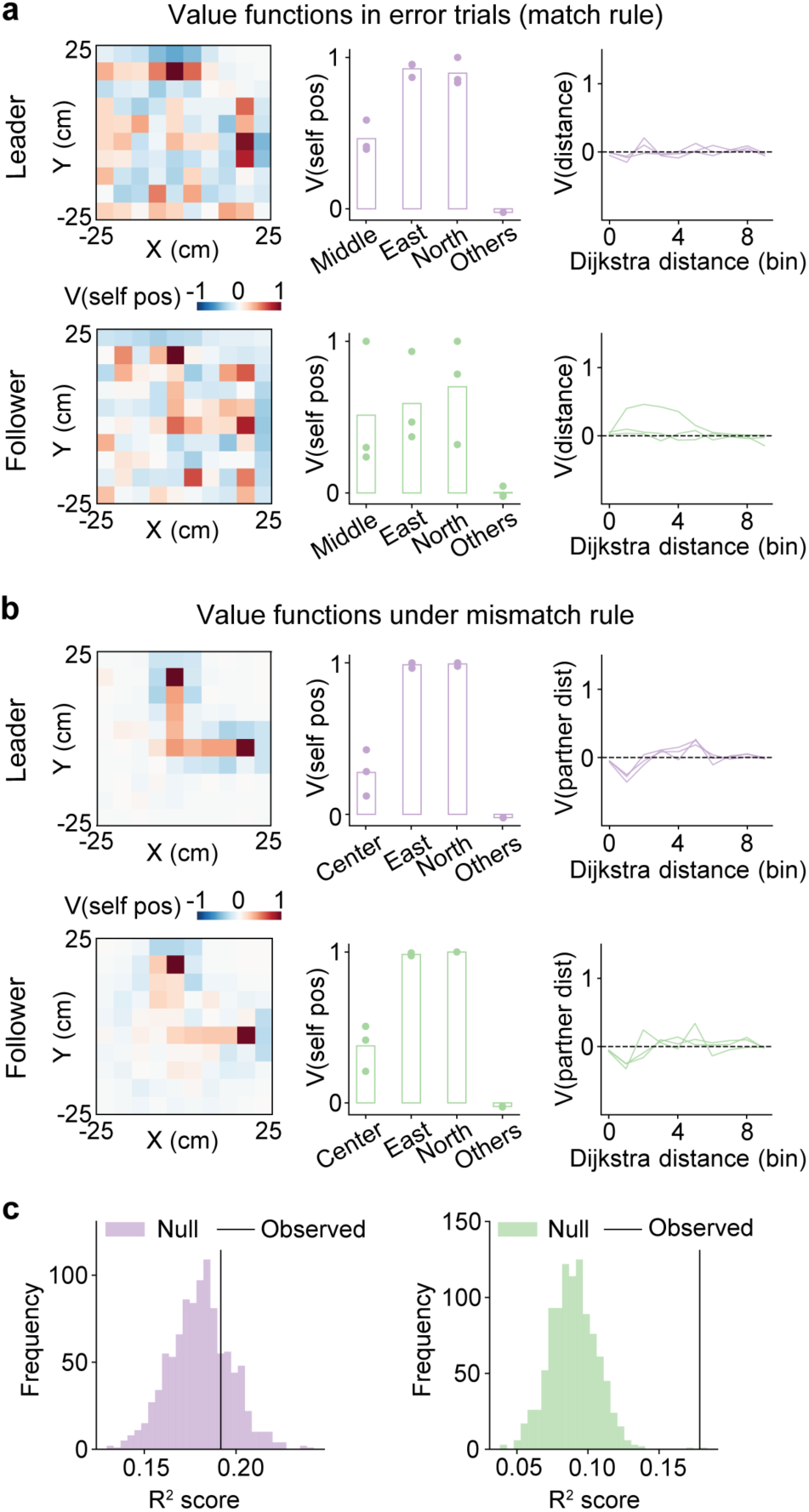
Context-dependent social value representations and their neural correlates. **a**, Inferred value functions in error trials for leaders (top row) and followers (bottom row); same as in Fig. 6e. **b**, Inferred value functions under the mismatch rule from leaders (top row) and followers (bottom row). Followers no longer assign higher value to proximal partner locations than leaders (paired *t*-test, *P* = 0.63; *n* = 3 animals). **c**, Coefficient of determination (R²) from linear regression predicting the MAIRL-inferred value function using population neural activity: leader (left, permutation test, *P* = 0.25), follower (right, \*\*\**P* = 3.0 × 10^-8^). Null distributions were generated by circularly shifting value labels 1,000 times; P-values reflect the probability of the observed R² under a Gaussian fit to the null.

## Methods

### Mice

Wild-type C57BL/6J female and male mice (2-6 months old; Jackson Laboratory) were used as experimental animals. All the mice were maintained on a 12/12-h reversed light/dark cycle (07:00 off, 19:00 on) with ad libitum access to food and water. During training and experiments, mice were water-restricted but maintained at 80-90% of their initial body weights. Experiments were conducted during the dark cycle, and each training session lasted for about 1 h, during which mice received 0.5-1.5 mL of water from the task. Animals received supplemental water as necessary to maintain their body weights. All procedures complied with the National Institutes of Health Guide for the Care and were approved by the Icahn School of Medicine at Mount Sinai Institutional Animal Care and Use Committee.

### Behavioral apparatus

The training arena was an 18 × 18-inch white-acrylic square chamber. Four reward zones were positioned at the center of each arena wall, with two adjacent water delivery ports. Reward ports were 3D-printed using white and transparent resins, incorporating an infrared beam for nose-poke detection and a white LED to indicate active ports. Water was delivered through a stainless-steel tube within each port, controlled by a solenoid valve (Lee Co., LHDB0533418H). An initiation detector, fitted with an infrared LED and a reflective sensor, was positioned at the center of the arena to detect trial initiation when approached by mice. Analog signals from all ports were digitized via Arduino Nano and acquired through a DAQ system (National Instruments, USB-6001). Behavioral control was implemented in LabVIEW to manage trial timing, LED signaling, water delivery, and data logging. Mouse behavior was recorded at 30 frames per second using an overhead camera (Teledyne FLIR, BFS-U3-16S2M-CS).

### Behavioral training

Prior to training, mice were water-restricted to 1 mL/day for at least three days and habituated to the experimenter and the behavioral setup. Training consisted of four main stages:

#### Stage 1: Light-guided water retrieval

Mice were trained to associate illuminated ports with water rewards within a single reward zone, separated from the rest of the arena by a transparent acrylic barrier. On each trial, either the left or right port was randomly illuminated; poking the illuminated port triggered water delivery and turned off the light. Mice advanced to Stage 2 after three days of training.

#### Stage 2: Learning to initiate trials

Mice were given access to the entire arena including all four reward zones and the central initiation spot. Trials began when a mouse is detected by the initiation spot, triggering illumination of both ports in a single active reward zone. Mice must poke an illuminated port within 15 s to receive a water reward; Poking an inactive port terminated the trial and was followed by a timeout (6-10 s). Mice reaching 80% correct trials (correct/total) progressed to Stage 3.

#### Stage 3: Social foraging shaping

Two cage mates (not necessarily siblings) that completed Stage 2 were introduced into the arena together. Either mouse could initiate a trial, illuminating both ports within a single reward zone. Both mice were required to poke the illuminated ports in the same zone within 15 s to receive a reward. Poking an inactive port by either mouse terminated the trial and triggered a timeout (6-10 s). Mice reaching 80% correct trials advanced to Stage 4a.

#### Stage 4a: Social foraging training

Following trial initiation, two of the four reward zones were randomly illuminated as active zones. Both mice were required to choose the same reward zone to receive a water reward. Trials were terminated without reward if the mice chose different reward zones (mismatch error) or if either mouse poked an inactive zone (unrewarded error). All error trials were followed by a timeout (6-10 s) added to the inter-trial interval (ITI, 3 s) before the next trial. Lapsed trials (failure to respond within 15 s in early Stage 4a or 8 s in late Stage 4a) were rare (0.014; Supplementary Fig. 1c) and excluded from correct rate calculations. Criterion performance was defined as 80% correct across three consecutive sessions.

#### Stage 2b: Solo foraging control

Solo foraging was performed typically after Stage 2 to assess the reward zone preferences of individual mice in the absence of a partner. Following initiation, two reward zones were randomly illuminated, and the mouse received water by poking any port from the active zones. Poking an inactive port ended the trial without a reward.

#### Stage 4d: Rule reversal

In Stage 4d, task contingency was reversed: mice were rewarded for choosing different reward zones. Choosing the same reward zone terminated the trial without reward. All other task parameters remained unchanged. A subset of mouse pairs well-trained in Stage 4 were retrained in Stage 4d.

### Video tracking

We used SLEAP, an open-source deep-learning-based framework for multi-animal pose tracking^64^, to estimate the mouse position and trajectory. To enable identification of individual mice, we shaved the back fur of one mouse in each pair. Each mouse was annotated with three key points: nose (tip of the nose), neck (center of the neck), and torso (center of mass), which were connected to form a skeleton. Approximately 4,200 video frames from ∼10 mouse pairs were manually annotated for training a SLEAP model, which was then used to automatically estimate the key points of other pairs. To optimize tracking accuracy, we additionally applied the Ensemble Kalman Smoother (EKS), a post-processing method that refines pose estimation outputs by smoothing several model predictions, resulting in more robust tracking^65^.

Custom MATLAB scripts (MathWorks, MATLAB 2024a) were used to align the behavior data from LabVIEW with the video frames based on the LED onset and offset. Reward zones were defined as semicircles with a 10 cm radius centered at the midpoint between each pair of reward ports. Mouse position was defined using its neck position, which was also used to calculate speed and acceleration.

### Tube test

To assess social hierarchy during the social foraging task, we conducted tube tests in a round-robin manner with four cage mates^66^. Each pair was tested multiple times throughout training, including Stage 2/2b, Stage 3, early Stage 4a, and well-trained (late) Stage 4a. Tests were conducted ∼30 minutes after social foraging sessions. Mice were placed at opposite ends of a 12-inch-long, 1-1/4-inch diameter transparent acrylic tube and encouraged to enter. Since only one mouse could pass through at a time, the one that forced the other out was defined as the dominant individual. Each group underwent six rounds of testing, and the experiment ended once a stable hierarchy was observed for four consecutive days.

### Social role assignment

Once a pair reached the training criterion (≥80% correct rate for three consecutive sessions), we assigned each mouse as either a leader or follower, and independently as an initiator or responder. To determine leadership, we computed the proportion of trials led by one mouse and compared it to chance (0.5) using a two-tailed binomial test (α = 0.01). If the distribution was significantly biased, the mouse that led more trials was designated the leader; otherwise, no leader was assigned. The same procedure was applied to assign initiators and responders. Importantly, these classifications reflect role biases over sessions rather than fixed identities— leaders may follow on some trials, and responders may initiate. Unrewarded errors and lapsed trials were excluded from analysis, as leader and follower roles could not be assigned in these trials, and both occurred infrequently (0.83% and 1.4%, respectively, during the well-trained stage; Supplementary Fig. 1b,c). Accordingly, we exclude unrewarded errors and lapses to evaluate mouse performance and define cooperation rate as the proportion of correct trials relative to the sum of correct and mismatch trials.

### Partner swapping

Four cage mates were first trained individually in the solo foraging task, then randomly assigned into two dyads for social foraging training. Once both pairs reached the training criterion and the social roles are assigned, we swapped partners either by pairing the leader from one dyad and the follower from the other (Fig. 1p,q), or by pairing both leaders or both followers (Fig. 1r). If both mice in a new pair were shaved, we marked one with black dye (Stoelting, 50450) for tracking with SLEAP. We then resumed training with the new pairs until they again reached the training criterion.

### Manual annotation of behavioral motifs

Trial start and end times were extracted from the behavioral recordings to segment the continuous video into individual trials. A 30-frame pre-trial buffer was appended to each segment to provide temporal context. Segmented clips were concatenated into a single video (5-15 minutes in duration) for manual annotation. Videos were then annotated frame by frame using Avidemux by trained observers. For each identified behavioral motif, the start and end frame numbers were recorded. We then mapped these indices to corresponding frames in the original video and generated Gantt charts for visual inspection. Motifs were quality-checked and excluded if they exceeded a predefined duration threshold or occurred during the ITI.

The following four social behavioral motifs were defined and manually annotated. All behavioral motifs reflect pairwise interactions, but the “track,” “sharp turn,” and “join” motifs were attributed to the mouse that played the more active role in the interaction. The “synchronized travel” motif was not assigned to either mouse, as it involved joint, symmetric behavior.

***Track***: One mouse, while initially approaching an active reward zone, slows down and turns its head to monitor the movement of its partner that is further away. Upon detecting the partner’s approach, it adjusts its timing to allow both animals arrive at the reward ports nearly simultaneously.

***Synchronized travel (Sync)***: The pair move in parallel, maintaining similar linear and angular velocities and a consistent close distance throughout the trajectory. They arrive at the same reward zone in a coordinated, synchronized manner.

***Sharp turn***: Each mouse initially moves toward a different active reward zone. During the approach, one mouse abruptly changes its trajectory by more than 90°, redirecting its movement toward the reward zone selected by the partner.

***Join***: One mouse initiates movement toward an active zone independently. The partner mouse, initially stationary or undecided, subsequently aligns its trajectory with the first mouse after observing its movement. Both animals then travel together toward the same reward zone.

### Virus

The following viral vectors were purchased from Addgene: AAV8-hSyn-hM4D(Gi)-mCherry (7 × 10^12^ vg/mL; 50475-AAV8), AAV8-hSyn-mCherry (1 × 10^13^vg/mL; 114472-AAV8), AAV9-syn-jGCaMP8m-WPRE (1 × 10^13^ vg/mL; 162375-AAV9), AAV9-Syn-GCaMP6f-WPRE-SV40 (7 × 10^12^ vg/mL; 100837-AAV9), and AAV5-CAG-GFP (7 × 10^12^ vg/mL; 37825-AAV5). AAV8-CaMKIIa-Kali1-eYFP (1.8 × 10^13^ vg/mL; GVVC-AAV-292) was obtained from the Stanford University Virus Core.

### Stereotaxic surgery

Mice were anesthetized with a cocktail of ketamine (100 mg/kg) and xylazine (10 mg/kg) and secured in a stereotaxic frame (Kopf Instruments, Model 940). Anesthesia was maintained with 1-1.5% isoflurane throughout the procedure. Viral injections were performed using a glass capillary connected to a nanoinjector (Drummond Scientific, Nanoject III) at a rate of 1-2 nL/s. Stereotaxic coordinates were determined according to the Paxinos and Franklin mouse brain atlas. Following surgery, mice received either a single dose of the long-acting analgesic buprenorphine (3.25 mg/kg, Ethiqa XR) or carprofen (5 mg/kg) for three consecutive days.

Chemogenetic inactivation of the mPFC was performed using AAV8-hSyn-hM4D(Gi)-mCherry, with pAAV8-hSyn-mCherry as the control. Viral injections were administered at 4 sites per hemisphere to cover the entire mPFC (Bregma coordinates: AP +1.98 mm, ML ±0.45 mm, DV - 1.85 mm/-1.45 mm, 200 nL per depth; AP +1.18 mm, ML ±0.45 mm, DV -1.10 mm, 300 nL; AP +0.50 mm, ML ±0.45 mm, DV -0.80 mm, 300 nL; AP -0.20 mm, ML ±0.45 mm, DV -0.80 mm, 300 nL). For chemogenetic inactivation of the OFC, 300 nL of AAV8-hSyn-hM4D(Gi)-mCherry was injected bilaterally at AP +2.46 mm, ML ±1.00 mm, DV -1.75 mm.

For optogenetic inactivation of mPFC, AAV8-CaMKIIa-Kali1-eYFP was used, with AAV5-CAG-GFP as the control. Viral injections were performed at two sites per hemisphere (300 nL per site): AP +1.98 mm, ML ±0.45 mm, DV -1.85 mm/-1.45 mm; and AP +1.18 mm, ML ±0.45 mm, DV -1.10 mm. Following injections, bilateral optical fibers (Amuza Inc., TeleLCD-Y-2.0-500-0.9) were implanted at AP +1.98 mm, ML ±0.45 mm, DV -0.80 mm. Optical fibers were secured with black dental acrylic (Lang Dental Manufacturing Co., Ortho-Jet™) to firmly attach the fiber and block external light.

To implant GRIN (Gradient Index) lens for miniscope recording, we performed a craniotomy over the mPFC at AP +1.98 mm, ML +0.45 mm with a 1-mm-radius window. Brain tissue above the mPFC was aspirated with a 27-gauge blunt or bent needle while continuously irrigating with cortex buffer to preserve tissue integrity. The resulting cavity was shaped to be as close to cylindrical as possible. 600 nL of AAV9-Syn-jGCaMP8m-WPRE or AAV9-Syn-GCaMP6f-WPRE-SV40 virus was then injected unilaterally at AP: +1.98 mm, ML: +0.45 mm, DV: -1.70 mm. After injection, a 1-mm-diameter, 4-mm-length GRIN lens (Inscopix) was positioned at the injection site and implanted 1.45 mm below the brain surface. The lens was secured with cyanoacrylate adhesive (Loctite) and further protected with a layer of low-toxicity silicone adhesive (World Precision Instruments, Kwik-Sil). Finally, dental acrylic was applied to stabilize the implant and cover the remaining exposed skull.

After 2-3 weeks of viral expression, mice were re-anesthetized and placed back in the stereotaxic frame. The overlying dental cement was carefully drilled off to expose the implanted GRIN lens. A Miniscope^67^ pre-mounted on a fixed baseplate, was positioned above the lens, and the field of view was monitored in real-time. The Miniscope was gradually lowered until imaging field was visible. Once we observed the optimal field of view, the baseplate was secured with cyanoacrylate and dental cement. To minimize external light interference, an outer layer of black dental cement was applied. A protective cap was then attached with screws, and mice were allowed to recover with postoperative analgesia.

### Chemogenetics

For chemogenetic manipulation, we injected the viral vectors into untrained mice and allowed at least 4 weeks of viral expression while simultaneously training them in the social foraging task. We administered the DREADD agonist Clozapine N-oxide (CNO; 5 mg/kg; Tocris No., 4936) dissolved in saline via intraperitoneal injection 30 minutes before the start of the session. We performed the control session the day before the inactivation session, when we injected mice with an equal volume of saline to control for potential effects of liquid intake and handling. To control for potential non-specific effects of CNO, we administered the same dose to a separate control group expressing mCherry and evaluated their behavior in the social foraging task.

### Wireless optogenetics

We used a wireless optogenetic system (Amuza Inc., Teleopto) for mPFC inactivation. After completing behavioral training, virus injection, and fiber implantation surgery, mice were given 2–3 weeks for viral expression and post-surgical recovery. Only mice that reached the training criterion were included in the optogenetic experiments, all conducted within four weeks post-surgery.

For stimulation, a lightweight receiver was connected to the implanted optic fibers. Stimulation signals were generated by the social foraging task control system (National Instruments, NI-6001) as TTL pulses and transmitted via BNC cables to the optogenetic control unit. The behavioral setup enabled precise inactivation across different task periods, including the entire trial duration and 1 s or 2 s from trial onset. During stimulation sessions, light pulses were delivered on one-third of the trials according to a pseudorandomized schedule. To prevent intensity fluctuations caused by battery discharge, we limited stimulation intensity to one-third of the maximum output of the implanted LED (590 nm, ∼0.8 mW per hemisphere). We fitted both animals with dummy receivers during three days of pre-experiment training to acclimate the mice to the wireless system. Optogenetic inhibition in both or a single animal was achieved by independently activating the receiver in each mouse.

### One-photon calcium imaging

Two post-surgical mice with clear imaging fields were paired and trained in the social foraging task. Calcium imaging was initiated upon reaching Phase 4a of the training protocol. Each day, one mouse underwent calcium imaging while the other was fitted with a dummy scope of equal weight to control for potential impact from the Miniscope attachment. Calcium imaging data were acquired using the Miniscope DAQ system and synchronized with behavioral video recordings. A TTL signal generated by Miniscope DAQ was transmitted via a trigger cable (6-pin GPIO Hirose Connector Cable, FLIR) to the behavior-recording camera, ensuring frame-by-frame alignment. Imaging was conducted at 30 frames per second, synchronized with video acquisition. We used Suite2p^68^ for ROI identification and extraction of calcium transients, and Cascade^69^ to infer spike rate from the calcium signals. To compute fluorescence changes (ΔF/F), we first subtracted 70% of the local neuropil signal from the fluorescence signal of each cell to obtain the raw trace. To estimate a dynamic baseline, we first smoothed the raw trace using a Gaussian filter with standard deviation σ = 6.7 s to reduce high-frequency noise and then applied a two-step filtering process: First, a moving minimum filter with a window size of 500 s was applied to the smoothed trace to track the lower envelope; then, a moving maximum filter of the same window size was applied to this result to obtain a conservative estimate of the baseline. ΔF/F was computed by normalizing the raw trace to this dynamic baseline.

### Electrophysiological recording and data analysis

Extracellular single-unit recordings were performed acutely in head-fixed animals using 64-channel silicon probes (Cambridge Neurotech, Acute H3 probe). Probes were stereotaxically targeted to the mPFC using Bregma coordinates: AP 1.78-2.22 mm, ML 0.45 mm, DV 1.85-2.30 mm. Four animals were used for chemogenetic validation and three animals for optogenetic validation.

In acute recording sessions, a craniotomy (0.3-0.8 mm in diameter) was performed over the target area prior to each session. Recording depth was determined using micromanipulator readings. After each session, the brain surface was protected with silicone gel (Dow Corning, 3-4680) and Kwik-Sil (World Precision Instruments). Recording sites were visualized by staining the probes with Vybrant DiO (Invitrogen, V22886) or Vybrant DiI (Invitrogen, V22885).

To validate inactivation using KALI-1, we positioned an optical fiber (RWD, 0.50 NA, 200 μm diameter) approximately 0.5 mm above the brain surface, producing a light spot with an estimated radius of ∼0.3 mm for delivering 590 nm illumination. To prevent collision with the fiber, the recording probe was inserted at a 10-degree angle. The brain surface was illuminated at varying light intensities, with each power level tested 10 times within a recording session (2 s of illumination per light pulse, 30 s between pulses).

For validation of the wireless optogenetic device, an optical fiber (Amuza inc.) was implanted at a 10-degree angle, secured 0.5 mm below the brain surface with dental cement. A craniotomy (0.3–0.8 mm diameter) was made at the implantation site. A recording probe was inserted at a 10-degree angle to avoid collision and positioned to record from mPFC neurons beneath the illuminated area. Recording sites were marked with DiI and verified histologically post-recording.

Recording data from the 64-channel probes was sampled at 25 kHz using the RHD recording system (Intan Inc.). Voltage signals were high-pass filtered and automatically sorted with Kilosort3^70^. Spike clusters were then manually curated using the Phy GUI^71^ to merge spikes from the same units and exclude noise and poorly isolated units.

### Behavioral analysis

To estimate how the decisions of the initiator are influenced by the partner, we compared the initiator’s zone choice in the cooperative foraging task to a solo foraging control. For this analysis, all adjacent trial types were rotated to align the active zones to a north-east configuration. Trials longer than 5 s (10.0% of all trials) and unrewarded errors and lapses (0.83% and 1.4%, respectively; Supplementary Fig. 1b,c) were excluded. For visualizing initiator choice (Fig. 2h,i; Supplementary Fig. 2b,c), we performed the following logistic regression:

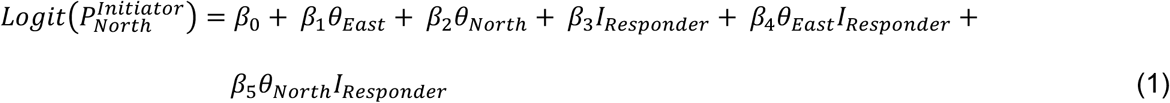

where 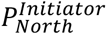 denotes the probability that the initiator selects the north zone. *θ_East_* and *θ_North_* are the initiator’s absolute head angles relative to the east and north directions, respectively*. I_responder_* indicates whether the responder is oriented toward a given reward zone (Fig. 2h,i) or whether the responder is located in a given reward zone (Supplementary Fig. 2b,c), depending on the condition. The *β* terms are fitted coefficients: *β*_0_ captures baseline bias toward the north zone on control trials (in log odds units); *β*_1_ and *β*_2_ reflect the animal’s sensitivity to heading information in control trials (with *β*_1_ > 0, *β*_2_ < 0 in well-trained animals); *β*_3_ quantifies the shift in bias due to the responder; and *β*_4_ and *β*_5_ reflect changes in heading sensitivity in the presence of the responder.

To estimate changes in heading sensitivity and directional bias induced by the presence of the responder (Fig. 2j,k; Supplementary Fig. 2d,e), we first computed the difference in head angles (Δθ, in degrees) relative to the east and north directions, reflecting the degree to which the heading is more aligned with the north direction than the east:

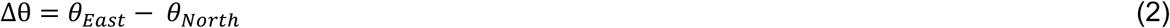

We then used Δθ to perform the following logistic regression:

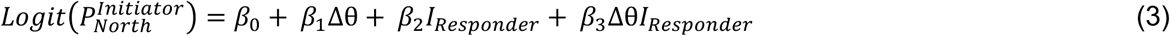

Similar to Equation (1), *β*_2_ quantifies the changes in choice bias due to the Responder; *β*_3_ estimates the changes in sensitivity to the initiator’s own heading in the presence of the responder.

To examine decision-making across all trials for both leaders and followers (Fig. 2r-t), we first calculated the differences in each animal’s heading and distance relative to the two active zones at trial onset, and standardized these values to the range of [0,1] prior to model fitting:

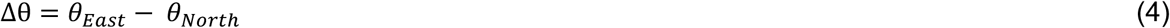

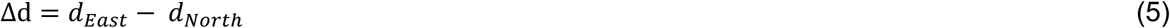

We then used the following equations to fit the leader and follower’s decision, respectively:

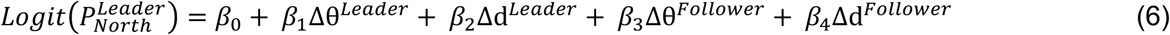

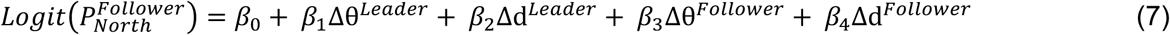

We predicted the leader and follower’s trial-by-trial choice using these logistic fits (Fig. 2s). The GLM was fit to a training set, and model performance was evaluated on a held-out test set as the proportion of correctly predicted choices. We used 90% of the data for training and 10% holdout for cross-validation, repeated over 100 folds. In each fold, the dataset was randomly split into the training and test sets. To assess statistical significance, we compared model performance against a shuffled null distribution. For each fold, a shuffled version of the data was created by randomly permuting choice labels, and the model was trained and tested using the same predictor variables. Model accuracy on true versus shuffled data was compared across folds using a bootstrap procedure (10,000 iterations) to estimate the P-value for the observed difference in mean accuracy. This procedure was applied separately to predict leader and follower choices, yielding role-specific estimates.

To examine the impact of chemogenetic and optogenetic inactivation on heading sensitivity and choice bias (Fig. 3i,m,q,v), we used the following logistic regression to fit the leader or follower’s decision:

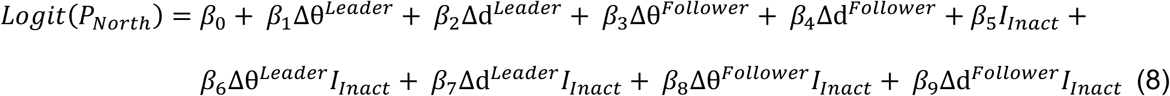

Where *β*_6_ to *β*_9_ quantify the changes in sensitivity to leader and follower’s heading and distance induced by inactivation. Trials were pooled across sessions and animals to increase statistical power.

### Choice selectivity analysis

To identify neurons selective for the animal’s reward zone choice (Fig. 4a-b), we computed mean calcium responses within the 1-s time window before trial end across trials for each neuron. We then used a one-way ANOVA to test for significant differences in response across the four choice conditions separately for each neuron. Neurons with a P-value below 0.05 were classified as significantly choice-selective.

To quantify the strength of choice tuning across neurons, we computed a variance-based selectivity index (SI) defined as:

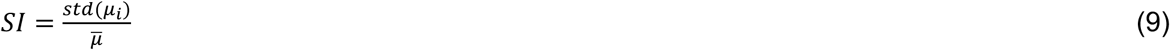

where *µ_i_* is the mean response of the neuron for each choice condition *i*, *std*(*µ*_i_)s the standard deviation across those means, and *µ̄* is the grand mean. This SI captures how much a neuron’s activity deviates across choice conditions relative to its overall activity level.

To assess whether neuronal population activity encoded zone choice, we trained a support vector machine (SVM) decoder to classify choice labels from trial-aligned calcium signals from each session. Neural activity was extracted from ΔF/F traces in a window spanning 2 s before and 2 s after trial end (corresponding to 60 pre-and 60 post-alignment frames). To reduce noise, we smoothed the traces by computing the mean responses in non-overlapping, 3-frame time bins. At each time bin, we trained a linear SVM (LIBLINEAR implementation^72^) to classify the animal’s choice for one of the four reward zones based on randomly sampled subsets of 100 simultaneously recorded neurons. Trials were randomly split stratified by choice into training (90%) and test (10%) sets across 50 cross-validation folds. Classification accuracy was computed as the mean correct rate on held-out trials. This procedure was repeated across time bins to obtain a temporal profile of decoding performance. All analysis was performed separately for leader and follower roles and for each session and then averaged across all available sessions from each animal.

### Leadership selectivity analysis

To quantify the evolution of leadership selectivity over time (Fig. 4h,i), we computed a trial-by-trial selectivity index (SI) for each neuron across the peri-arrival window. ΔF/F traces were aligned to the arrival of the recorded mouse, and a fixed time window (−2 s to +3 s, corresponding to 60 pre- and 90 post-alignment frames) was extracted for analysis. Trials in which leadership could not be determined due to ties in arrival times were excluded. For each time point and neuron, we calculated an area under the receiver operating characteristic curve (auROC) comparing activity on trials where the recorded animal led vs. followed. The auROC was linearly scaled to the range of [-1, 1] by computing SI = 2 × (auROC - 0.5), yielding a time-by-neuron matrix of SI values. To visualize population dynamics, we plotted the SI heatmap across neurons sorted by the timing of their peak selectivity. Positive SI values indicate stronger activity when leading, and negative values indicate greater activity when following.

To identify neurons selective for trial-by-trial leadership identity (Fig. 4o), we computed mean ΔF/F responses within a 1s time window following the arrival of the recorded mouse for each trial. Trials in which leadership could not be determined due to ties were also excluded. For each neuron, responses were grouped by whether the recorded animal was leading or following on a trial-by-trial basis. We then quantified leadership selectivity using the auROC-based SI as described above. To assess statistical significance, we generated a null distribution of SI values using stratified label shuffling. Specifically, to control for potential spatial choice confounds, we permuted the leadership labels within each port choice category (left vs. right) independently.

This procedure was repeated 1,000 times per neuron, and a P-value was calculated as the fraction of shuffled SIs whose absolute magnitude exceeded that of the observed SI. Neurons with P < 0.05 were classified as significantly selective for leadership identity.

To further examine whether mPFC population activity encodes trial-by-trial leadership dynamics (Fig. 4p), we trained a logistic regression decoder to classify whether the recorded animal led or followed on each trial. We excluded trials with ties in arrival times and aligned calcium traces to time of arrival. For each neuron, ΔF/F signals were binned into non-overlapping windows (3-frame bins). For each time bin, we trained a logistic regression model (LIBLINEAR, L2-regularized) on 90% of trials and tested on the remaining 10%, stratified by leadership labels. If a test set lacked representation from either class, we incrementally increased the holdout proportion until both classes were included. This procedure was repeated over 50 cross-validation folds. Decoding performance was quantified as the mean auROC across all test folds.

To establish significance, we compared decoding performance against a null distribution generated from random permutations of leadership labels. For each permutation, we applied the same decoding pipeline and recorded the corresponding auROC.

### Spatial selectivity analysis

To identify neurons selective for allocentric or egocentric positions of the self and the partner (Fig. 6a-g, Supplementary Fig. 5a-c), we computed deconvolved spike rate maps in four spatial reference frames: (1) the position of the recorded mouse in the arena (allocentric self), (2) the partner’s position in the arena (allocentric partner), (3) the partner’s position in egocentric coordinates without heading alignment (egocentric partner unaligned with HD), and (4) the partner’s position aligned to the heading of the recorded mouse (egocentric partner aligned with HD). All maps were constructed using 5 x 5 cm spatial bins. We included all frames in a session but excluded the frames when either mouse is within any reward zone (10 cm from any reward ports) to rule out reward-related confounds. For each neuron, we quantified spatial selectivity using three criteria. Only neurons passing all three criteria in a given reference frame were classified as selective for that spatial variable.

#### Spatial information content

Spatial information was calculated^73^ as:

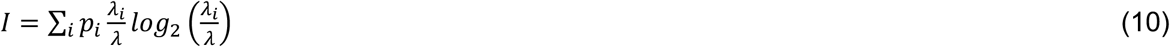

where *λ_i_* is the mean spike rate in the *i^th^* spatial bin, *λ* is the overall mean spike rate, and *pi* is the occupancy probability of the *i^th^* bin. Significance was assessed via circularly shifting the deconvolved spike rate (n = 100 permutations) relative to the position labels. Cells with spatial information exceeding the 95th percentile of the null distribution were considered significant.

#### Spatial coherence

We computed the mean correlation between each spatial bin and the mean of its 8 neighbors, across all bins. Cells with coherence values above the 95th percentile of a circularly shifted null distribution were deemed coherent.

#### Within-session stability

To assess tuning stability within a session, we computed the Spearman correlation between rate maps generated from the first and second halves of the session. Neurons were considered consistent if their correlation exceeded the 95th percentile of a null distribution obtained by circularly shifting spike times.

### Decoding partner distance and egocentric angle

To decode partner distance and egocentric angle on a frame-by-frame basis (Fig. 6h-k), we applied logistic regression classifiers to the population activity. The partner distance was discretized into 5-cm bins spanning 0-35 cm, and egocentric partner angle was divided into 15 uniform bins (-180° to 180°, positive when partner is on the left). Neurons from multiple imaging sessions of the same animals were pooled after binning. To balance class sizes, an equal number of frames were randomly sampled without replacement from each bin across sessions. Sampled frames were then sorted by their original timestamps to preserve their temporal order for decoding. Spike rates from all included neurons were extracted for each frame, and the corresponding distance or angle bin index was used as the class label.

To assess decoding accuracy, we implemented multi-class logistic regression using the LIBLINEAR solver (L2-regularized). A five-fold cross-validation was used, in which the data were evenly split into five non-overlapping segments. In each fold, four segments were used for training and the remaining segment for testing, such that all data segments were used once as the test set. Decoding was repeated 10 times with 300 randomly selected neurons per repetition.

For each cross-validation fold and repetition, we recorded the predicted class probabilities for test frames and averaged them across repetitions to form a confusion matrix, representing the probability of predicting each distance or angle bin given the ground truth. As a performance metric, we computed the proportion of predictions matching the true distance label (Fig. 5i), or the proportion of predictions falling within ±1 bin of the true angle label, circularly defined (Fig. 5k). To assess statistical significance, final decoding accuracy was compared to a null distribution obtained by decoding shuffled data generated through circularly shifting the deconvolved spike traces. A confusion matrix was also generated for the shuffled data (Supplementary Fig. 6a,b).

### CEBRA-Behavior multi-session embedding

We trained each CEBRA embedding on a single behavioral feature (e.g., partner distance or egocentric angle), using 80% of trials pooled across all sessions from a single animal for training, and evaluated model performance on the remaining 20% of held-out trials. To assess statistical significance, each embedding was compared to a temporally shuffled control, in which the behavioral variable was shifted by a random but identical time offset across both training and test sets. CEBRA embeddings were configured with a batch size of 500, a constant temperature of 0.01, a hidden layer size of 32, an output dimensionality of 3, a learning rate of 0.0003, and a cosine distance metric. The delta conditional distribution was used, and training was run for 1,000 iterations using the offset10-model architecture.

To quantify the structure of CEBRA embeddings, we discretized the behavior variable of interest into 5 quintiles containing an equal number of points. We then defined a modified silhouette score for each data point *i* as:

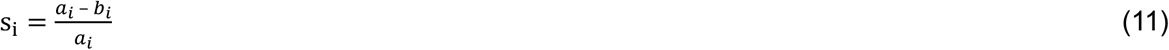

where *a_i_* is the mean distance between point *i* and all points in different quintiles, *b_i_* is the mean distance between point *i* and all other points within the same quintile. This is the standard medoid silhouette score except that the maximum in the denominator is replaced with a mean, increasing sensitivity to structured but non-clustered data.

### Partner distance and egocentric angle tuning

To characterize neuronal tuning to the egocentric location of a conspecific, we constructed tuning curves for partner distance and angle using calcium imaging data collected across animals and sessions (Fig. 5p,q,s,t). Neurons were first selected based on significant tuning to egocentric partner position, identified via 2D egocentric spatial maps as described above (Fig. 5e). For distance tuning, only neurons exhibiting significant spatial information for partner distance (*P* < 0.05 vs. circularly shifted data) were included. For partner angle tuning, only neurons with a significant Rayleigh vector length (*P* < 0.05 vs. circularly shifted data) were retained. Data were pooled across multiple imaging sessions and animals.

For each selected neuron, the tuning curve was normalized to its peak firing rate to allow comparison across neurons. The resulting matrices of normalized tuning curves were sorted by each neuron’s peak bin location (distance or egocentric angle) and visualized as heatmaps. The population distribution of peak distances or angles was summarized using histograms.

### Multi-agent reinforcement learning (MARL) modeling

We developed a forward MARL model to simulate cooperative behavior in a spatial foraging task similar to the mouse paradigm. The environment was discretized into an 11 × 11 square grid in which two agents, represented by their spatial coordinates, learn to navigate toward and jointly occupy randomly activated reward zones. The global state at each time step was defined as:

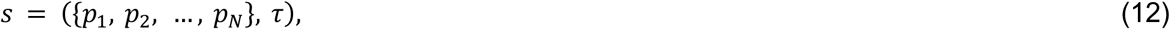

where 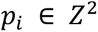 denotes the spatial coordinates of the agent *i*, and τ encodes the task phase (pre-activation vs post-activation) and the pair of active reward locations. N = 2 agents in this simulation.

Each agent *i* independently learns a state-value function *Q_i_*(*s*), which was updated using temporal-difference learning:

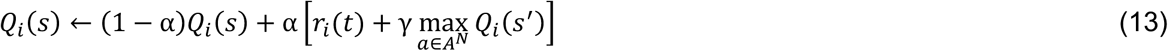

where *α* is the learning rate, γ is the discount factor, and *r_i_*(*t*) is the reward received at time *t*. Action selection followed an epsilon-greedy policy over the *joint* action set *A^N^*, where *A* = {*stay, up, down, left, right*}. With probability ϵ, a uniformly random action was selected; otherwise, agents evaluated possible joint actions and selected the move leading to the state *s’* that maximized *Q_i_*(*s’*). Invalid moves (off-grid) were masked out.

Trials began with agents randomly placed on the grid. Training consisted of two stages: (1) initiation, where agents navigated toward the center of the arena to activate rewards; and (2) cooperative foraging, where both agents were trained to simultaneously arrive at the same active reward zone. Successful coordination yielded a reward for each agent; penalties were applied for entering inactive zones (unrewarded error), choosing different active targets (mismatch error), or exceeding a maximum trial duration without reward (lapse). Each movement incurred a small energy cost. Training continued until convergence, with environment resets upon reward collection, errors, or timeouts.

### Multi-agent inverse reinforcement learning (MAIRL) modeling

#### Pre-processing of foraging trajectories

To pre-process the foraging trajectories, each mouse was represented as a massless point anchored at its neck position. The arena was discretized into a 10 ×10 grid with 5-cm spatial bins and each animal’s position was assigned to the nearest grid point. To minimize artifacts introduced by discretizing the arena, we excluded time steps where both animals remained stationary. Each animal had nine possible actions: stay, up, down, left, right, up-left, up-right, down-left, and down-right.

#### Model setup

We modeled the cooperative foraging task as a Markov Decision Process (MDP) involving two agents with shared objectives. This MDP is defined by the tuple 〈𝒮, 𝒜, 𝒫, 𝑟, 𝛾〉, where 𝒮 = 𝒮_1_ × 𝒮_1_ represents the joint environmental state formed as the Cartesian product of individual state sets 𝒮_i_; In our task, each agent occupies one of the 100 discrete spatial locations, yielding 10,000 possible joint states. 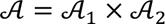. denotes the set of joint actions, with each agent having 9 valid options, resulting in 81 total joint actions. 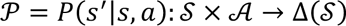 defines the transition dynamics, where Δ represents a probability distribution over 𝒮. We assumed deterministic transitions such that a specific joint action 𝑎 always leads to a predetermined next state *s’.* The reward function 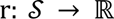 maps the current joint state 𝑠 to a scalar value. γ ∈ [0,1] is the discount factor for future rewards.

We then estimated the value map that maximizes the likelihood of the observed trajectory. Following the inverse reinforcement learning (IRL) framework^74–76^, given 〈𝒮, 𝒜, 𝒫, 𝛾〉 and 𝑁 trials of both agents’ trajectories 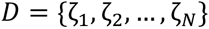, we infer the unknown reward functions 𝑟 such that P(D|r) is maximized. Each trajectory ζ*_i_* consists of independent joint state-action pairs 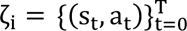. Consequently, the posterior probability of observing expert trajectory 𝜁_i_ can be calculated under the specific policy π derived from value function 𝑟, as shown in the following equation:

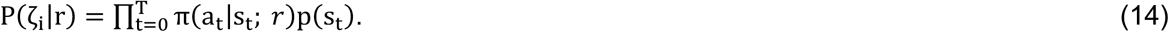

Therefore, the core of this maximization problem is the parameterization of the joint policy function 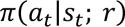, using a probabilistic modeling of social interactions based on multi-agent reinforcement learning (MARL). Note that policy function π could be derived from value function 𝑣 using the value iteration algorithm, and both share the same underlying parameterization.

#### Parameterization of joint value function

To infer the reward function 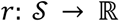 in the two-agent foraging task, we must estimate |𝒮| = 10000 parameters. This is computationally demanding in a multi-agent setting, as the size of the joint state space |𝒮| grows exponentially with the number of agents. To address this, we implemented a value decomposition approach, allowing the joint value function to be expressed as a sum of marginal and interaction components, as defined by:

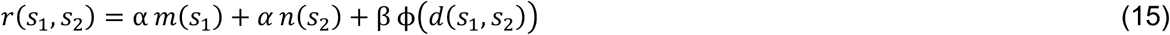

where 𝑠_i_ is agent 𝑖’s current location, 𝑑 denotes the Dijkstra distance between 𝑠_1_ and 𝑠_2_(i.e. the shortest number of steps required to travel between two locations), and α, β are linear coefficients describing the weights of the value map functions. This approach drastically reduces the number of parameters from 10,000 to 212. Numerical analyses and experiments^46^ demonstrated that this value decomposition achieves negligible reconstruction error relative to the original joint value function. Consequently, our objective then is to learn 𝛼, *β*, 𝑚, 𝑛, 𝜙 to maximize 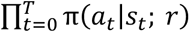 over the observed state-action pairs {𝑠_$_, 𝑎_$_}.

#### Maximum entropy policy formulation

To facilitate the inference problem, we adopted a differentiable maximum entropy policy formulation to derive the policy from the joint value function^75^

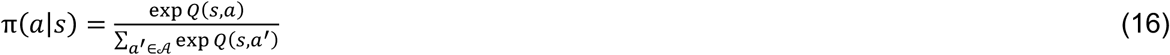

where Q(𝑠, 𝑎) is a soft Q-function obtained by performing soft value iteration:

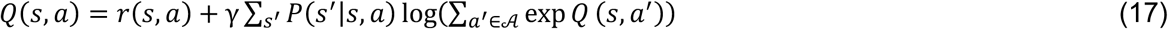

Note that a temperature term is not needed in the SoftMax function, as it is absorbed into the absolute value of 𝑄 and subsequently 𝑟, which is the variable we aim to estimate.

#### Inference algorithm

The inference procedure is detailed previously^48^. Our objective is to learn 𝛼, *β*, 𝑚, 𝑛, 𝜙 to maximize 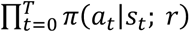 over 𝑁 observed trials. We assume all parameters follow a Gaussian prior with known variance and zero mean, with the weight coefficients following 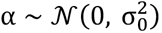, and 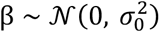. Incorporating priors to the map functions 𝑚, 𝑛, 𝜙 is equivalent to adding an L-2 regularizer with coefficients λ_1_ and 𝜆_2_ to their entries. Mathematically, the parameter optimization progress could be written as:

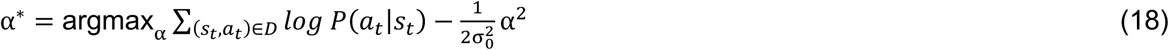

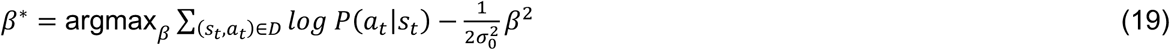

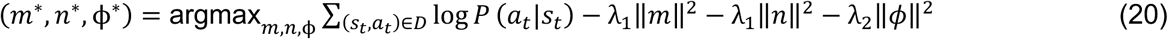

We used coordinate ascent to iteratively update the weights 𝛼, *β* and the maps 𝑚, 𝑛, 𝜙 while holding the other parameters fixed. Separate learning rates η_1_ and η_2_ were used for updating the weights and the maps, respectively.

We used 80% of the trials for model fitting and the remaining 20% to calculate the test log-likelihood as model evidence. Three random searches were initialized with different seeds and the one with the highest test log-likelihood was used. The preset hyperparameters were learning rates η_1_ = 0.1, η_2_ = 0.005 and regularization strengths λ_1_ = 5 , λ_2_ = 1, selected for interpretability of the recovered value functions of the physical locations (i.e. 𝑚(𝑠_1_) and 𝑛(𝑠_2_)).

Without appropriate regularization, the recovered value maps tend to exhibit randomly activated regions. The exact values of the regularization parameters were not critical, as long as they imposed sufficient sparsity constraints. All other hyperparameters were searched from the range shown in Table 1. Based on test set log-likelihood and convergence speed, we chose γ = 0.90, 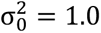.

**Table 1.** Hyperparameter search ranges.

| Parameter | Range |
| --- | --- |
| $\gamma$ | [0.70,0.90,0.99] |
| $\sigma_0^2$ | [0.02,0.2,1.0] |

#### Estimation of individual trajectories

In the above sections, we maximize the likelihood of observing the joint action pair, 𝑃(𝑎|𝑠) = 𝑃(𝑎_1_, 𝑎_2_|𝑠_1_, 𝑠_2_), which incorporates information from both agents. Given the asymmetric behavior observed in the cooperative foraging task, it is also informative to estimate the individual value functions that govern each animal’s decision-making. In this section, subscripts refer to individual agents, and the time index *t* is omitted for clarity. Assuming that animals have independent control over their decisions, we separate the probability into

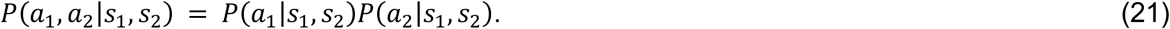

This formulation allows us to model the decision-making process of an individual animal. Without losing generality, we aimed to identify the reward function 𝑟^1^(𝑠_1_, 𝑠_2_) for animal 1 such that *P*(*a*_1_|*s*_1_,*s*_2_;*r*^1^) is maximized. Following the policy derivation described in the earlier section, this reward function 𝑟^1^ defines a joint policy *π*(*a*_1_,*a*_2_|*s*_1_,*s*_2_;*r*^1^), from which we derived the marginal policy:

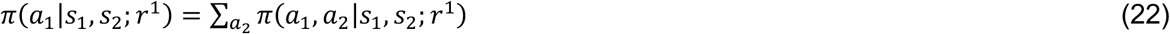

The remaining steps in the inference procedure followed accordingly, using the marginalized policy in place of the joint policy.

#### Incorporating egocentric angle information

The partner’s egocentric angle θ was calculated as described in the previous sections. The angle was then discretized into the front (-90° to 90°) or the back (90° to 180° and -180° to -90°) fields. Consequently, the discretized 𝜃 takes the value of either 0 or 1. Angle information was provided to the Markov chain as input. We therefore maximized 𝑃(𝑎|𝑠, 𝜃) given 𝑟(𝑠|θ). To further simplify the dependence on angle, we allowed the decomposed joint value function to depend on θ only through the interaction term:

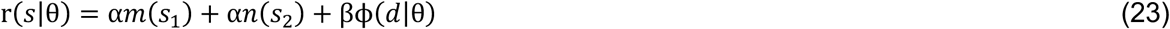

#### Model selection and comparison

We focused on different parameterizations of the decomposed joint value function for model selection (Fig. 6; Table 2). These models were nested in complexity. For each model, we estimated its fit upon convergence of the inference procedure described in the *Inference algorithm* section. Model comparisons between nested models were conducted using a chi-square test, with degrees of freedom equal to the difference in the number of parameters. The baseline model (model with the allocentric position of self) was additionally compared to a constant prediction model, whose log-likelihood was computed as the number of decision pairs times 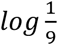.

**Table 2.**
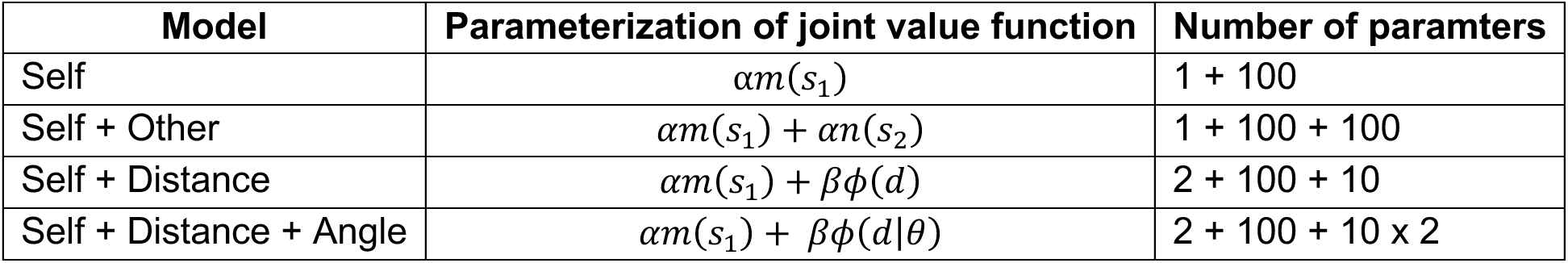
Model formulation under different hypotheses.

#### Simulation of foraging trajectory to predict reward zone choice

To intuitively assess model fitting performance, we simulated foraging trajectories using the inferred joint value function and predicted the reward zone choice by the animals. To simulate the trajectory for an animal in each trial, we initialized the animal at the same starting position as in the observed data. At each time step, the animal’s action was sampled from its marginalized policy *π*(*a*_1_|*s*_1_,*s*_2_;*r*^1^), derived from the inferred joint value function. The selected action determined the animal’s next location, while its partner location was taken directly from the observed trajectory. This allowed us to simulate individual decisions in a two-agent foraging setup. The simulation is terminated when the animal reaches the reward zone. The prediction is considered correct if the simulated choice matches the observed behavior. This process was repeated for all the trials in each session, and the correct prediction rate was calculated for each model. Note that this procedure was not applied to models incorporating angle information, as angle dynamics could not be simulated in this setup.

#### Decoding of inferred value from population neural activity

To determine whether mPFC neural activity encodes estimated value functions from our model, we performed linear decoding of value estimates 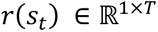 from population activity *N_t_* ∈ ℝ^NxT^, on a session-by-session basis. The dataset was randomly split into 80% time steps for training and 20% time steps for test. Model performance was quantified using the *R*^2^ score on the test set. To assess statistical significance, we generated a null distribution by applying circular time shifts to the neural data and repeating the decoding procedure 1,000 times (Supplementary Fig. 7b). The P-value was calculated by fitting a Gaussian distribution to the null 𝑅. scores and computing the percentile rank of the observed 𝑅. within this distribution. The minimum P-value was capped at 10^−16^ to account for numerical precision.

### Use of LLM

A large language model (ChatGPT, OpenAI) was used to assist with language editing and refinement of the manuscript text. All scientific content, interpretations, and conclusions were developed by the authors.

## Data availability

Data will be made available at publication.

## Code availability

Code will be made available at publication.

## Acknowledgements

We thank K. Deisseroth for the AAV-CaMKIIa-KALI1-eYFP construct; L. Paninski for advice on behavioral analysis; S.J. Russo, D. Cai, T. Shuman, B. Sweis, H. Morishita, X. Wu, Z. Pennington, A. Baggetta, Y. Zaki, Z. Wick, P. Philipsberg, F.K. Chiang, L. Li, R.D.-d. Cuttoli, and H. Li for advice on experiments; W. Janssen, B. Wu, E. Sullaway, C. Shamsu, and N. Tillison for assistance; and P.J. Kenny, P. Rudebeck, M.N. Shadlen, and C. Dulac for comments on the manuscript. This work was supported by the Alfred P. Sloan Foundation, the NIMH (MH133039), the Chan-Zuckerberg Initiative, the Simons Foundation, the Esther A. & Joseph Klingenstein Fund, and the Friedman Brain Institute (Z.H.W.); a Seaver Fellowship (Y. Cheng); a Swartz Postdoctoral Fellowship (Y. Chen); the Air Force Office of Scientific Research (FA9550-22-1-0337) (N.R.); and the NIMH (MH132653) (T.P.).

## Author contributions

H.Z.W. conceptualized and supervised the project. H.Z.W., Y. Cheng, Y. Chen, and N.Y. designed the experiments. Y. Cheng collected the data with assistance from M.K., R.L., T.S., and A.K. Y. Chen developed the MAIRL algorithm. H.Z.W., Y. Cheng, and Y. Chen analyzed the data with inputs from E.S.S. and N.R. R.P.K. and E.S.S. performed the CEBRA analysis. R.S., J.W., N.R., and Y. Chen performed the MARL modeling. U.K. contributed to developing the behavioral task and analysis. T.P. and M.W. advised on behavioral analysis. H.Z.W. wrote the manuscript with inputs from all authors.

## Competing interests

H.Z.W. is an inventor on a patent applied for by the Icahn School of Medicine at Mount Sinai that covers the cooperative foraging paradigm described here.

## References

1 Wilson, E. O. Sociobiology : the new synthesis. (Cambridge, Mass. : Belknap Press of Harvard University Press, 1975., 1975).

2 Krause, J. & Ruxton, G. D. Living in Groups. (Oxford University Press, 2002).

3 Axelrod, R. & Hamilton, W. D. The Evolution of Cooperation. Science 211, 1390–1396 (1981). 10.1126/science.7466396

4 Nowak, M. A. Five rules for the evolution of cooperation. Science 314, 1560–1563 (2006). 10.1126/science.1133755

5 King, A. J., Johnson, D. D. & Van Vugt, M. The origins and evolution of leadership. Curr Biol 19, R911–916 (2009). 10.1016/j.cub.2009.07.027

6 Nagy, M., Akos, Z., Biro, D. & Vicsek, T. Hierarchical group dynamics in pigeon flocks. Nature 464, 890–893 (2010). 10.1038/nature08891

7 Van Vugt, M., Hogan, R. & Kaiser, R. B. Leadership, followership, and evolution: some lessons from the past. Am Psychol 63, 182–196 (2008). 10.1037/0003-066X.63.3.182

8 Uhl-Bien, M., Riggio, R. E., Lowe, K. B. & Carsten, M. K. Followership theory: A review and research agenda. The Leadership Quarterly 25, 83–104 (2014). 10.1016/j.leaqua.2013.11.007

9 Luo, L., Callaway, E. M. & Svoboda, K. Genetic Dissection of Neural Circuits: A Decade of Progress. Neuron 98, 256–281 (2018). 10.1016/j.neuron.2018.03.040

10 Carandini, M. & Churchland, A. K. Probing perceptual decisions in rodents. Nat Neurosci 16, 824–831 (2013). 10.1038/nn.3410

11 Rilling, J. K. & Sanfey, A. G. The neuroscience of social decision-making. Annu Rev Psychol 62, 23–48 (2011). 10.1146/annurev.psych.121208.131647

12 Wang, H. & Kwan, A. C. Competitive and cooperative games for probing the neural basis of social decision-making in animals. Neurosci Biobehav Rev 149, 105158 (2023). 10.1016/j.neubiorev.2023.105158

13 von Neumann, J., Morgenstern, O. & Rubinstein, A. Theory of Games and Economic Behavior (60th Anniversary Commemorative Edition). (Princeton University Press, 1944).

14 Yu, J. H., Napoli, J. L. & Lovett-Barron, M. Understanding collective behavior through neurobiology. Curr Opin Neurobiol 86, 102866 (2024). 10.1016/j.conb.2024.102866

15 Erlich, J. C. et al. Mice dynamically adapt to opponents in competitive multi-player games. bioRxiv, 2025.2002.2014.638359 (2025). 10.1101/2025.02.14.638359

16 Gold, J. I. & Shadlen, M. N. The neural basis of decision making. Annu Rev Neurosci 30, 535–574 (2007). 10.1146/annurev.neuro.29.051605.113038

17 Hanks, T. D. et al. Distinct relationships of parietal and prefrontal cortices to evidence accumulation. Nature 520, 220–223 (2015). 10.1038/nature14066

18 Khilkevich, A. et al. Brain-wide dynamics linking sensation to action during decision-making. Nature 634, 890–900 (2024). 10.1038/s41586-024-07908-w

19 Newsome, W. T., Britten, K. H. & Movshon, J. A. Neuronal correlates of a perceptual decision. Nature 341, 52–54 (1989). 10.1038/341052a0

20 Heekeren, H. R., Marrett, S. & Ungerleider, L. G. The neural systems that mediate human perceptual decision making. Nat Rev Neurosci 9, 467–479 (2008). 10.1038/nrn2374

21. Platt, M. L. & Glimcher, P. W. Neural correlates of decision variables in parietal cortex. Nature 400, 233–238 (1999). 10.1038/22268

22 Padoa-Schioppa, C. & Assad, J. A. Neurons in the orbitofrontal cortex encode economic value. Nature 441, 223–226 (2006). 10.1038/nature04676

23 Montague, P. R., Dayan, P. & Sejnowski, T. J. A framework for mesencephalic dopamine systems based on predictive Hebbian learning. J Neurosci 16, 1936–1947 (1996). 10.1523/JNEUROSCI.16-05-01936.1996

24 Rangel, A., Camerer, C. & Montague, P. R. A framework for studying the neurobiology of value-based decision making. Nat Rev Neurosci 9, 545–556 (2008). 10.1038/nrn2357

25 Schultz, W. Predictive reward signal of dopamine neurons. J Neurophysiol 80, 1–27 (1998). 10.1152/jn.1998.80.1.1

26 Wittmann, M. K. et al. Causal manipulation of self-other mergence in the dorsomedial prefrontal cortex. Neuron 109, 2353–2361 e2311 (2021). 10.1016/j.neuron.2021.05.027

27 Forli, A. & Yartsev, M. M. Hippocampal representation during collective spatial behaviour in bats. Nature 621, 796–803 (2023). 10.1038/s41586-023-06478-7

28 Testard, C. et al. Neural signatures of natural behaviour in socializing macaques. Nature 628, 381–390 (2024). 10.1038/s41586-024-07178-6

29 Ray, S. et al. Hippocampal coding of identity, sex, hierarchy, and affiliation in a social group of wild fruit bats. Science 387, eadk9385 (2025). 10.1126/science.adk9385

30 Zada, D. et al. Development of neural circuits for social motion perception in schooling fish. Curr Biol 34, 3380–3391.e3385 (2024). 10.1016/j.cub.2024.06.049

31 Arora, S. & Doshi, P. A survey of inverse reinforcement learning: Challenges, methods and progress. Artificial Intelligence 297, 103500 (2021). 10.1016/j.artint.2021.103500

32 Ashwood, Z. C., Jha, A. & Pillow, J. W. in *Proceedings of the 36th International Conference on Neural Information Processing Systems* Article 2151 (Curran Associates Inc., New Orleans, LA, USA, 2022).

33 Pereira, T. D. et al. SLEAP: A deep learning system for multi-animal pose tracking. Nat Methods 19, 486–495 (2022). 10.1038/s41592-022-01426-1

34 Biderman, D. et al. Lightning Pose: improved animal pose estimation via semi-supervised learning, Bayesian ensembling and cloud-native open-source tools. Nat Methods 21, 1316–1328 (2024). 10.1038/s41592-024-02319-1

35 Lin, D. et al. Functional identification of an aggression locus in the mouse hypothalamus. Nature 470, 221–226 (2011). 10.1038/nature09736

36 Li, S. W. et al. Frontal neurons driving competitive behaviour and ecology of social groups. Nature 603, 661–666 (2022). 10.1038/s41586-021-04000-5

37 Padilla-Coreano, N. et al. Cortical ensembles orchestrate social competition through hypothalamic outputs. Nature 603, 667–671 (2022). 10.1038/s41586-022-04507-5

38 Wang, F. et al. Bidirectional control of social hierarchy by synaptic efficacy in medial prefrontal cortex. Science 334, 693–697 (2011). 10.1126/science.1209951

39 Amodio, D. M. & Frith, C. D. Meeting of minds: the medial frontal cortex and social cognition. Nat Rev Neurosci 7, 268–277 (2006). 10.1038/nrn1884

40 Apps, M. A., Rushworth, M. F. & Chang, S. W. The Anterior Cingulate Gyrus and Social Cognition: Tracking the Motivation of Others. Neuron 90, 692–707 (2016). 10.1016/j.neuron.2016.04.018

41 Yizhar, O. et al. Neocortical excitation/inhibition balance in information processing and social dysfunction. Nature 477, 171–178 (2011). 10.1038/nature10360

42 Yamamuro, K. et al. A prefrontal-paraventricular thalamus circuit requires juvenile social experience to regulate adult sociability in mice. Nat Neurosci 23, 1240–1252 (2020). 10.1038/s41593-020-0695-6

43 Kingsbury, L. et al. Correlated Neural Activity and Encoding of Behavior across Brains of Socially Interacting Animals. Cell 178, 429–446 e416 (2019). 10.1016/j.cell.2019.05.022

44 Wang, P. Y. et al. Transient and Persistent Representations of Odor Value in Prefrontal Cortex. Neuron 108, 209–224 e206 (2020). 10.1016/j.neuron.2020.07.033

45 Wilson, R. C., Takahashi, Y. K., Schoenbaum, G. & Niv, Y. Orbitofrontal Cortex as a Cognitive Map of Task Space. Neuron 81, 267–279 (2014). 10.1016/j.neuron.2013.11.005

46 Jiang, M. et al. Evolution and neural representation of mammalian cooperative behavior. Cell Rep 37, 110029 (2021). 10.1016/j.celrep.2021.110029

47 Schneider, S., Lee, J. H. & Mathis, M. W. Learnable latent embeddings for joint behavioural and neural analysis. Nature 617, 360–368 (2023). 10.1038/s41586-023-06031-6

48 Chen, Y., Radulescu, A. & Wu, H. Z. Unveiling the latent dynamics in social cognition with multi-agent inverse reinforcement learning. bioRxiv, 2024.2010.2009.617461 (2024). 10.1101/2024.10.09.617461

49 Rudebeck, P. H., Buckley, M. J., Walton, M. E. & Rushworth, M. F. A role for the macaque anterior cingulate gyrus in social valuation. Science 313, 1310–1312 (2006). 10.1126/science.1128197

50 Chang, S. W., Gariépy, J. F. & Platt, M. L. Neuronal reference frames for social decisions in primate frontal cortex. Nat Neurosci 16, 243–250 (2013). 10.1038/nn.3287

51 Haroush, K. & Williams, Z. M. Neuronal prediction of opponent’s behavior during cooperative social interchange in primates. Cell 160, 1233–1245 (2015). 10.1016/j.cell.2015.01.045

52 Ong, W. S., Madlon-Kay, S. & Platt, M. L. Neuronal correlates of strategic cooperation in monkeys. Nat Neurosci 24, 116–128 (2021). 10.1038/s41593-020-00746-9

53 Zhang, K. M. et al. A new paradigm of learned cooperation reveals extensive social coordination and specific cortical activation in mice. Mol Brain 16, 40 (2023). 10.1186/s13041-023-01032-y

54 Rose, M. C., Styr, B., Schmid, T. A., Elie, J. E. & Yartsev, M. M. Cortical representation of group social communication in bats. Science 374, eaba9584 (2021). 10.1126/science.aba9584

55 Witkowski, P. P., Park, S. A. & Boorman, E. D. Neural mechanisms of credit assignment for inferred relationships in a structured world. Neuron 110, 2680–2690.e2689 (2022). 10.1016/j.neuron.2022.05.021

56 Danjo, T., Toyoizumi, T. & Fujisawa, S. Spatial representations of self and other in the hippocampus. Science 359, 213–218 (2018). 10.1126/science.aao3898

57 Omer, D. B., Maimon, S. R., Las, L. & Ulanovsky, N. Social place-cells in the bat hippocampus. Science 359, 218–224 (2018). 10.1126/science.aao3474

58 Zhang, X. et al. Multiplexed representation of others in the hippocampal CA1 subfield of female mice. Nature Communications 15, 3702 (2024). 10.1038/s41467-024-47453-8

59 Murugan, M. et al. Combined Social and Spatial Coding in a Descending Projection from the Prefrontal Cortex. Cell 171, 1663–1677 e1616 (2017). 10.1016/j.cell.2017.11.002

60 Frith, U. & Frith, C. D. Development and neurophysiology of mentalizing. Philos Trans R Soc Lond B Biol Sci 358, 459–473 (2003). 10.1098/rstb.2002.1218

61 Dunbar, R. I. M. The social brain hypothesis. *Evolutionary Anthropology: Issues*, News, and Reviews 6, 178–190 (1998). 10.1002/(SICI)1520-6505(1998)6:5<178::AID-EVAN5>3.0.CO;2-8

62 Lockwood, P. L., Apps, M. A. J. & Chang, S. W. C. Is There a ’Social’ Brain? Implementations and Algorithms. Trends Cogn Sci 24, 802–813 (2020). 10.1016/j.tics.2020.06.011

63 Wittmann, M. K. et al. Basis functions for complex social decisions in dorsomedial frontal cortex. Nature (2025). 10.1038/s41586-025-08705-9

## Method references

64 Pereira, T. D. et al. Fast animal pose estimation using deep neural networks. Nature Methods 16, 117–125 (2019). 10.1038/s41592-018-0234-5

65 Biderman, D. et al. Lightning Pose: improved animal pose estimation via semi-supervised learning, Bayesian ensembling and cloud-native open-source tools. Nature Methods 21, 1316–1328 (2024). 10.1038/s41592-024-02319-1

66 Fan, Z. et al. Using the tube test to measure social hierarchy in mice. Nat Protoc 14, 819–831 (2019). 10.1038/s41596-018-0116-4

67 Cai, D. J. et al. A shared neural ensemble links distinct contextual memories encoded close in time. Nature 534, 115–118 (2016). 10.1038/nature17955

68 Pachitariu, M. et al. Suite2p: beyond 10,000 neurons with standard two-photon microscopy. bioRxiv, 061507 (2017). 10.1101/061507

69 Rupprecht, P. et al. A database and deep learning toolbox for noise-optimized, generalized spike inference from calcium imaging. Nature Neuroscience 24, 1324–1337 (2021). 10.1038/s41593-021-00895-5

70 Pachitariu, M., Sridhar, S., Pennington, J. & Stringer, C. Spike sorting with Kilosort4. Nature Methods 21, 914–921 (2024). 10.1038/s41592-024-02232-7

71 Rossant, C. et al. Spike sorting for large, dense electrode arrays. Nature Neuroscience 19, 634–641 (2016). 10.1038/nn.4268

72 Fan, R.-E., Chang, K.-W., Hsieh, C.-J., Wang, X.-R. & Lin, C.-J. LIBLINEAR: A library for large linear classification. the Journal of machine Learning research 9, 1871–1874 (2008).

73 Skaggs, W., Mcnaughton, B., Gothard, K. & Markus, E. An information-theoretic approach to deciphering the hippocampal code. Advances in neural information processing systems 5, 1030–1037 (1993).

74. Abbeel, P. & Ng, A. Y. in *Proceedings of the twenty-first international conference on Machine learning*. 1.

75. Ziebart, B. D., Maas, A. L., Bagnell, J. A. & Dey, A. K. in *Aaai*. 1433-1438 (Chicago, IL, USA).

76 Ashwood, Z., Jha, A. & Pillow, J. W. Dynamic inverse reinforcement learning for characterizing animal behavior. Advances in neural information processing systems 35, 29663–29676 (2022).

